# Oral Glucosamine Ameliorates Aggravated Neurological Phenotype in Mucopolysaccharidosis III Type C Mouse Model Expressing Misfolded HGSNAT Variant

**DOI:** 10.1101/2021.08.26.457793

**Authors:** Xuefang Pan, Mahsa Taherzadeh, Poulomee Bose, Rachel Heon-Roberts, Annie L. A. Nguyen, TianMeng Xu, Camila Pará, Yojiro Yamanaka, David A. Priestman, Frances M. Platt, Shaukat Khan, Nidhi Fnu, Shunji Tomatsu, Carlos R. Morales, Alexey V. Pshezhetsky

**Affiliations:** CHU Sainte-Justine Research Center, University of Montreal, Montreal, H3T 1C5, QC, Canada; Department of Anatomy and Cell Biology, McGill University, Montreal, H3A 0C7, QC, Canada; Goodman Cancer Research Centre, McGill University, Montreal, H3A 1A3, QC, Canada; Department of Pharmacology, University of Oxford, Oxford UK; Nemours/Alfred I. duPont Hospital for Children, Wilmington, DE, USA

**Keywords:** Glycosaminoglycans, Mucopolysaccharidosis, Heparan sulfate acetyl-Coenzyme A: α-glucosaminide N-acetyltransferase, Knock-in Mouse Model, Molecular Chaperones.

## Abstract

**Objective:** Over 55% of mucopolysaccharidosis IIIC (MPS IIIC) patients have at least one allelic missense variant responsible for misfolding of heparan sulfate acetyl-CoA:*α*-glucosaminide N-acetyltransferase (HGSNAT). These variants are potentially treatable with pharmacological chaperones, such as a competitive HGSNAT inhibitor, glucosamine. Since the constitutive HGSNAT knockout mice, we generated previously cannot be used to test such strategy *in vivo,* we generated a novel model, the *Hgsnat^P304L^* strain, expressing misfolded mutant HGSNAT with human missense mutation Pro311Leu (Pro304Leu in the mouse enzyme).

**Results:** *Hgsnat^P304L^* mice present deficits in short-term (novel object recognition test) and working/spatial (Y-maze test) memory at 4 months of age, 2-4 months earlier than previously described gene-targeted *Hgsnat-Geo* mice, which lack HGSNAT protein. *Hgsnat^P304L^* mice also show increased severity of synaptic deficits in CA1 neurons, and accelerated course of CNS pathology including neuronal storage of heparan sulfate, accumulation of misfolded proteins, increase of simple gangliosides, and neuroinflammation as compared with *Hgsnat-Geo* mice. Expression of misfolded human Pro311Leu HGSNAT protein in cultured hippocampal *Hgsnat-Geo* neurons aggravated reduction of synaptic proteins. Memory deficits and majority of pathological changes in the brain were rescued in mice receiving daily doses of oral glucosamine.

**Interpretation:** Altogether, our data for the first time demonstrate dominant-negative effects of the misfolded HGSNAT Pro304Leu variant and show that these effects are treatable by oral administration of glucosamine, suggesting that patients, affected with missense mutations preventing normal folding of the enzyme, could benefit from chaperone therapy.

## Introduction

Mucopolysaccharidosis IIIC (MPS IIIC) or Sanfilippo syndrome type C is a rare genetic neurogenerative disease manifesting with neuropsychiatric problems, such as hyperactivity, aggressiveness and autistic features, followed by developmental delay, childhood dementia, speech and hearing loss [1]. Most patients become paraplegic during adolescence and die before adulthood, but some survive until the fourth decade of life [1] with progressive dementia and retinitis pigmentosa [2–5].

The disease is caused by deleterious variants in the gene encoding the lysosomal membrane enzyme, heparan sulfate acetyl-CoA: α-glucosaminide N-acetyltransferase (HGSNAT, EC 2.3.1.78), which catalyzes transmembrane acetylation of glucosamine residues of heparan sulfate (HS) before their hydrolysis by α-N-acetylglucosaminidase. Lysosomal storage of undegraded HS in the brain cells leads to neuroinflammation and neuronal dysfunction followed by neuronal death (reviewed in [6]). Of over seventy disease-causing *HGSNAT* variants, that have been identified in MPS IIIC patients, 35 are missense [7]. Expression studies showed that the resulting amino acid substitutions lead to the synthesis of misfolded HGSNAT protein, unable to escape the endoplasmic reticulum (ER) quality control and reach the lysosomes [8, 9]. These mutations are among the most frequent, with about 55% of MPS IIIC patients affected with at least one of them. Previously, we could partially rescue 10 mutant misfolded HGSNAT variants by treating patient’s cells with the inhibitor of HGSNAT, glucosamine [8], suggesting that these patients could potentially benefit from a pharmacological chaperone therapy (PCT), applicable to genetic disorders (including LSDs), where mutations result in the synthesis of proteins with amino acid substitutions.

Lysosomal enzymes are synthesized on polyribosomes and secreted into the lumen of the ER in a largely unfolded state [10]. The WT enzyme folds into the appropriate (native) conformation with the assistance of various molecular chaperones, such as BiP, heat shock proteins, calnexin, and calreticulin [11]. In contrast, although mutant enzymes might be stable and catalytically active in the acidic milieu of the lysosome, they are often not folded properly and cannot be transported to the lysosomes. These mutant proteins are retained in the ER and degraded by the proteosome- associated pathway [12]. Since folding is a thermodynamically controlled process, mutant proteins tend to fold into an alternative (misfolded) conformation because of their altered amino acid sequences. However, molecules, that mimic substrate binding in the active site, such as competitive inhibitors, may work as pharmacological chaperones (PC), stabilizing the proper position of active site residues and shifting the equilibrium toward the correctly folded state of the enzyme [13–19]. As a result, the correctly folded mutant enzyme passes the quality-control system of the ER and undergoes further maturation and normal transport to the lysosome. Once a mutant enzyme-chaperone complex reaches the lysosome, the chaperone is replaced by a highly concentrated (accumulated) substrate to allow the enzyme to function, leading to an increase in residual enzyme activity in the cell. Previous studies identified effective chaperones for several lysosomal enzymes, some restoring impaired catabolic pathways in cultured patient cells or showing therapeutic effect in mouse models of GM1 and GM2 gangliosidoses, Gaucher or Fabry diseases, reviewed elsewhere [20, 21]. After proof-of-concept studies, PCT is now being translated into clinical applications for cystic fibrosis [22] and for the lysosomal diseases, Fabry, Gaucher and Pompe [23–26]. PCT drug for the Fabry disease, Galafold [25], has received approval in the European Union and in the USA.

Since the constitutive knockout *Hgsnat-Geo* mice, we generated previously [27], cannot be used to test chaperone therapy *in vivo,* in the current study, we have produced the mouse line expressing HGSNAT with the human pathogenic variant Pro311Leu (Pro304Leu in the mouse protein). The mice homozygous for the *Hgsnat^P304L^* allele show a drastically bigger increase of blood HS level, lysosomal storage, and inflammatory cytokines and synaptic defects as compared with the knockout mice of similar age and have 2 months earlier onset of memory impairment consistent with the dominant-negative effect of the misfolded HGSNAT mutant. Behavioural problems, synaptic defects and majority of brain pathology were rescued by treating mice with daily doses of oral glucosamine validating the use of chaperone therapy as a promising approach to treat the MPS IIIC patients affected with missense mutations.

## Materials and Methods

The data, analytic methods, and study materials will be made available to other researchers for purposes of reproducing the results or replicating the procedure.

### Murine models

Approval for animal experimentation was granted by the Animal Care and Use Committee of the Ste-Justine Hospital Research Center. Mice were housed in an enriched environment with continuous access to food and water, under constant temperature and humidity, on a 12 h light/dark cycle. Mice were kept on a normal chow diet (5% fat, 57% carbohydrate). *Hgsnat-Geo* mice have been previously described [27]. *Hgsnat^P304L^* knock-in C57Bl/6J mouse strain was generated at McGill Integrated Core for Animal Modeling (MICAM) using CRISPR/Cas9 technology, targeting exon 9 of the *Mus musculus* heparan sulfate acetyl-CoA: alpha-glucosaminide N-acetyltransferase (*Hgsnat*) gene. To generate knock-in founders, a single guide nucleotide RNA

(sgRNA) was designed, using the CRISPR online tool (http://crispr.mit.edu), to target a genomic site on the murine *Hgsnat* locus with minimal potential off-target effects. sgRNA and Cas9 mRNA were microinjected into zygotes with single-stranded oligodeoxynucleotide (ssODN), barring a c.911C ˃T mutation encoding for the P304L change (**Supplementary Figure S1**, 100 ng/μl of ssODN, 20 ng/μl of sgRNA and 20 ng/μl of Cas9).

The zygotes were further cultured overnight in EmbryoMax KSOM drops (Millipore) covered with mineral oil (Irvine Scientific) in a 35 mm dish at 37°C in a 5% CO2 incubator. The embryos which developed up to the 2-cell stage were transferred to oviducts of pseudo-pregnant females to generate chimeric mice litter. Pups were delivered at full term and, at weaning, were genotyped by Sanger sequencing of single allele fragments, obtained by PCR amplification of genomic DNA from the tail clips. The adult founder mice containing the allele with the desired mutation were mated with C57BL/6N mice (Envigo, former Harlan Laboratories). Genotypes of the offspring were determined by amplifying a 988-bp product from purified mouse tail DNA (forward primer: ATGGAGTGCCTGATGGGAGG; reverse primer: GATCTAGAAACGGCCCGAAGA). Since the c.911C ˃T mutation eliminated the *NcoI* restriction site, to distinguish between mutant and WT amplicons, the PCR products were further digested with *NcoI* (New England Biolabs, Cat# R3193S) at 37°C for 1 h and then, run on a 2% agarose gel. Fragments of 688- and 300-bp were detected for the WT allele, and an undigested 988-bp fragment for the targeted *Hgsnat^P304L^* allele. To test for the presence of potential off-target effects, the sgRNA sequence (TGAACGTCGCCAGGTCGCGG) was blasted against the *Mus musculus* genome. Fragments of 763-bp of the *Spg7* gene exon 7, presenting the highest identity score, were amplified by PCR from genomic DNA of F1 and WT mice (forward primer: CTTTTGGAGCTGTGGCCCTA; reverse primer: AGTGGAAGATTTTGCCAGGC), and analyzed by Sanger sequencing for the presence of deleterious variants. This analysis did not reveal mutations in the *Spg7* gene in the F1 generation (**Supplementary Figure S2**). The F1 mice were bred to C57BL/6J mice to generate heterozygous mice that were further intercrossed to generate WT, heterozygote, and homozygous mutants.

### Enzyme activity assays

The specific enzymatic activities of HGSNAT, β-hexosaminidase, and β-galactosidase were assayed essentially as previously described (19). Tissues extracted from mice or pellets were snap-frozen in liquid nitrogen before storage at -80°C. Fifty mg samples were homogenized in 250 µl of H2O using a sonic homogenizer (Artek Systems Corporation). Cultured MEF cells were harvested by scraping and lysed in H2O by 10-sec sonication on ice (60 W). For HGSNAT assays, 5 µl aliquots of the homogenates were combined with 5 µl of McIlvain Buffer (pH 5.5), 5 µl of 3 mM 4-methylumbelliferyl-β-D-glucosaminide (Moscerdam), 5 µl of 5 mM acetyl-coenzyme A and 5 µl of H2O. The reaction was incubated for 3 h at 37°C, stopped with 975 µl of 0.4 M glycine buffer (pH 10.4), and fluorescence was measured using a ClarioStar plate reader (BMG Labtech). Blank samples were incubated without the homogenates which were added after the glycine buffer. The activity of β-hexosaminidase was measured by combining 2.5 µl of 10x diluted homogenate (∼ 2.5 ng of protein) with 15 µl of 0.1 M sodium acetate buffer (pH 4.2), and 12.5 µl of 3 mM 4- methylumbelliferyl N-acetyl-β-D-glucosaminide (Sigma-Aldrich) followed by incubation for 30 min at 37°C. The reaction was stopped 0.4 M glycine buffer (pH 10.4) and fluorescence was measured as above.

The activity of acidic β-galactosidase was measured by adding 12.5 µl of 0.4 M sodium acetate, 0.2 M NaCl (pH 4.2) and 12.5 µl of 1.5 mM 4-methylumbelliferyl β-D-galactoside (Sigma-Aldrich) to 10 µl of 10x diluted homogenate (∼1 ng of protein). After 15-min incubation at 37°C, the reaction was stopped with 0.4 M glycine buffer (pH 10.4) and fluorescence was measured as above.

HGSNAT N-acetyltransferase activity against BODIPY-glucosamine substrate in brain homogenates was performed by combining 6 µL of homogenate (∼6 µg protein), 6 µL of McIlvain’s phosphate/citrate buffer, pH 6.5, 4 µL of 10 mM acetyl CoA in H2O, and 4 µL of 40 mM BODIPY-glucosamine (1[4,4-difluoro-5,7-dimethyl-4-bora-3a,4a-diaza-s-indacene-3- propionyl-glycylamino]-*β*-D-glucosamine) [28]. After incubation for 3 h at 37°C in a 96-well PCR plates (BioScience Inc.), the reaction was terminated by the addition of 180 µL of 100 mM HCl. Twenty microliter aliquots of the final reaction were transferred to 96-well filter plate (Millipore, 40 mm nylon mesh), pre-embedded with 100 µL Toyopearl cation exchange media SP 650M (Tosoh) for each well. Prior to the assay, the resin was washed twice with 250 µL of H2O per well, and the plates were centrifuged at 50 g for 30 s to remove any excess water. The fluorescent neutral reaction product was eluted with four 90 µL aliquots of 1 M HCl by centrifugation of the plates at 50 g for 30 s. Combined eluent (360 µL) was transferred to 96-well Reader Black polystyrene plates (Life Science), and the amount of fluorescent product was measured.

### Behavioral analysis

The spontaneous alternation behavior, spatial working memory and exploratory activity of mice were evaluated using a white Y- maze as previously described [29]. The maze consisted of three identical white Plexiglas arms (40 × 10 × 20 cm, 120° apart) under dim lighting conditions. Each mouse was placed at the end of one arm, facing the center, and allowed to explore the maze for 8 min. All experiments were performed at the same time of the day and by the same investigator to avoid circadian and handling bias. Sessions were video-recorded and arm entries were scored by a trained observer, unaware of the mouse genotype or treatment. Successful alternation was defined as consecutive entries into a new arm before returning to the two previously visited arms. Alternation was calculated as: [number of alternations/total number of arm entries − 2] × 100.

Novel object recognition test was used for assessing short-term recognition memory [30, 31]. Mice were placed individually in a 44 × 33 × 20 cm (length x width x height) testing chamber with white Plexiglas walls for 10 min habituation period and returned to their home cage. The next day, mice were placed in the testing chamber for 10 min with two identical objects (red plastic towers, 3 x 1.5 x 4.5 cm), returned to the home cages, and 1 hour later, placed back into the testing chamber in the presence of one of the original objects and one novel object (a blue plastic base, 4.5 x 4.5 x 2 cm) for 10 min. After each mouse, the test arena as well as the plastic objects were cleaned with 70% ethanol to avoid olfactory cue bias. The discrimination index (DI) was calculated as the difference of the exploration time between the novel and old object divided by total exploration time. A preference for the novel object was defined as a DI significantly higher than 0 [32]. Mice who showed a side preference, noted as a DI of ± 0.20 during familiarization period, and those who had a total exploration times lower than 3 seconds were excluded from analysis.

The open-field test was performed as previously described [33]. Briefly, mice were habituated in the experimental room for 45 mins before the commencement of the test. Each mouse was then gently placed in the center of the open-field arena and allowed to explore for 20 min. The mouse was removed and transferred to its home cage after the test, and the arena was cleaned with 70% ethanol before the commencement of the next test. Analysis of the behavioral activity was done using the Smart video tracking software (v3.0, Panlab Harvard Apparatus), and total distance traveled and percent of time spent in the center zone were measured for hyperactivity and anxiety assessment, respectively.

The elevated plus-maze test was performed as described by Amegandjin, *et al* [33]. Each mouse was placed in the center of the elevated plus maze and allowed to freely explore undisturbed for 10 min. After each testing, the mouse was returned to the home cage and the arena was cleaned with 70% ethanol before the commencement of the next test. Analysis of the behavioral activity (percentage of time spent in the center zone, closed arms, and open arms; as well as the number of open arm entries) was done by the Smart v3.0 software.

### Transmission electron microscopy

At 3 and 6 months, 3 mice from each group were anesthetized with sodium pentobarbital (50 mg/kg BW) and perfused with PBS, followed by 2.5% glutaraldehyde in 0.2 M phosphate buffer (pH 7.2). The brains were extracted and post-fixed in the same fixative for 24 h at 4 °C. The hippocampi were dissected, mounted on glass slides, stained with toluidine blue and examined on a Leica DMS light microscope to select the CA1 region of the hippocampus for electron microscopy. The blocks were further embedded in Epon, and 100 nm ultrathin sections were cut with an Ultracut E ultramicrotome, mounted on 200-mesh copper grids, stained with uranyl acetate (Electron Microscopy Sciences) and lead citrate, and examined on a FEI Tecnai 12 transmission electron microscope. For quantification, the micrographs were taken with 13,000 x and 30,000 x magnification.

### Neuronal cultures

Primary hippocampal neurons were cultured from the brains of embryo at gestational day 16 (E16). The hippocampi were isolated and treated with 2.5% trypsin solution (Sigma-Aldrich, T4674) for 15 min at 37°C. The cells were washed 3 times with Hank’s Balanced Salt Solution (HBSS, Gibco, 14025-092) and mechanically dissociated by pipetting, using glass Pasteur pipettes with 3 different opening sizes (3, 2 and 1 mm). Then, they were counted with the viability dye trypan blue (ThermoFisher Scientific, 15250061), using a hemocytometer, and resuspended in Neurobasal media (Gibco, 21103-049) containing L-glutamine, B27, N2, penicillin and streptomycin. The cells were plated at a density of 60,000 cells per well, respectively, in a 12-well plate on coverslips previously coated with Poly-L-Lysine (Sigma Aldrich, P9155). Cells were cultured for 21 days, and 50% of media was changed every three days.

### Whole cell recordings in acute hippocampal slices

Acute hippocampal slices were prepared as described [34]. Briefly, animals were anaesthetized deeply with isoflurane and decapitated. The brain was dissected carefully and transferred rapidly into an ice-cold (0–4°C) solution containing the following (in mM): 250 sucrose, 2 KCl, 1.25 NaH2PO4, 26 NaHCO3, 7 MgSO4, 0.5 CaCl2 and 10 glucose, pH 7.4. The solution was oxygenated continuously with 95% O2 and 5% CO2, 330–340 mOsm/L. Transverse hippocampal slices (thickness, 300 µm) were cut using a vibratome (VT1000S; Leica Microsystems ), transferred to a bath at room temperature (23°C) with standard ACSF at pH 7.4 containing the following (in mM): 126 NaCl, 3 KCl, 1 NaH2PO4, 25 NaHCO3, 2 MgSO4, 2 CaCl2, 10 glucose, continuously saturated with 95% O2 and 5% CO2 and allowed to recover for 1 h. During the experiments, slices were transferred to the recording chamber at physiological temperature (30–33 °C) continuously perfused with standard ACSF, as described above, at 2 ml/min. Pyramidal CA1 neurons from the hippocampus were identified visually using a 40X water immersion objective. Whole-cell patch-clamp recordings were obtained from single cells in voltage- or current-clamp mode and only 1 cell per slice was recorded to enable post-hoc identification and immunohistochemical processing. Recording pipettes (4–6 MΩ) were filled with a K-gluconate based solution for voltage-clamp recordings (in mM): 130 K-gluconate, 10 KCl, 5 diNa-phosphocreatine, 10 HEPES, 2.5 MgCl2, 0.5CaCl2,1 EGTA, 3 ATP-Tris, 0.4 GTP-Li, 0.3% biocytin, pH 7.2–7.4, 280–290 mOsm/L.

After obtaining whole cell configuration, passive membrane properties were monitored for 5 min and current clamp recordings were done to measure action potential characteristics. Slices were then perfused with 0.5 μM TTX (to isolate miniature events) for 3 mins before commencing voltage clamp recordings. Cells were voltage clamped at -70 mV for mEPSCs recording and then held at 0 mV (calculated from the reversal potential of Cl) for mIPSCs recording. Data acquisition (filtered at 2–3 kHz and digitized at 15 kHz; Digidata 1440A, Molecular Devices, CA, USA) was performed using the Axopatch 200B amplifier and the Clampex 10.6 software (Molecular Devices). Both mEPSCs and mIPSCs were recorded for 7 min and a running template on a stable baseline (minimum of 30 events) was used for the analysis of miniature events on MiniAnalysis. Clampfit 10.2 software was used for analysis of action potential characteristics and other passive membrane properties.

For some experiments, to verify that all mEPSCs are blocked, slices were perfused with 5 μM DNQX (6,7-dinitroquinoxaline-2,3-dione) and 50 μM AP5 in addition to the TTX while recording mEPSCs at -70 mV after addition of α-amino-3-hydroxy-5-methyl-4-isoxazolepropionic acid (AMPA) and N-methyl-D-aspartate receptor (NMDAR) blockers. Similarly, for some experiments, slices were perfused with 100 μM BMI (bicuculline methiodide) and 50 μM AP5 in addition to the TTX in the ACSF to verify that all mIPSCS are blocked at 0 mV.

### Real-time qPCR

RNA was isolated from snap-frozen brain, kidney and liver tissues using the TRIzol reagent (Invitrogen) and reverse-transcribed using the iScript™ Reverse Transcription Supermix (Bio RAD #1708840) according to the manufacturer’s protocol. qPCR was performed using a LightCycler® 96 Instrument (Roche) and SsoFast™ EvaGreen® Supermix with Low ROX (Bio RAD #1725211) according to the manufacturer’s protocol. RLP32 mRNA was used as a reference control. The following forward (F) and reverse (R) primers have been used:

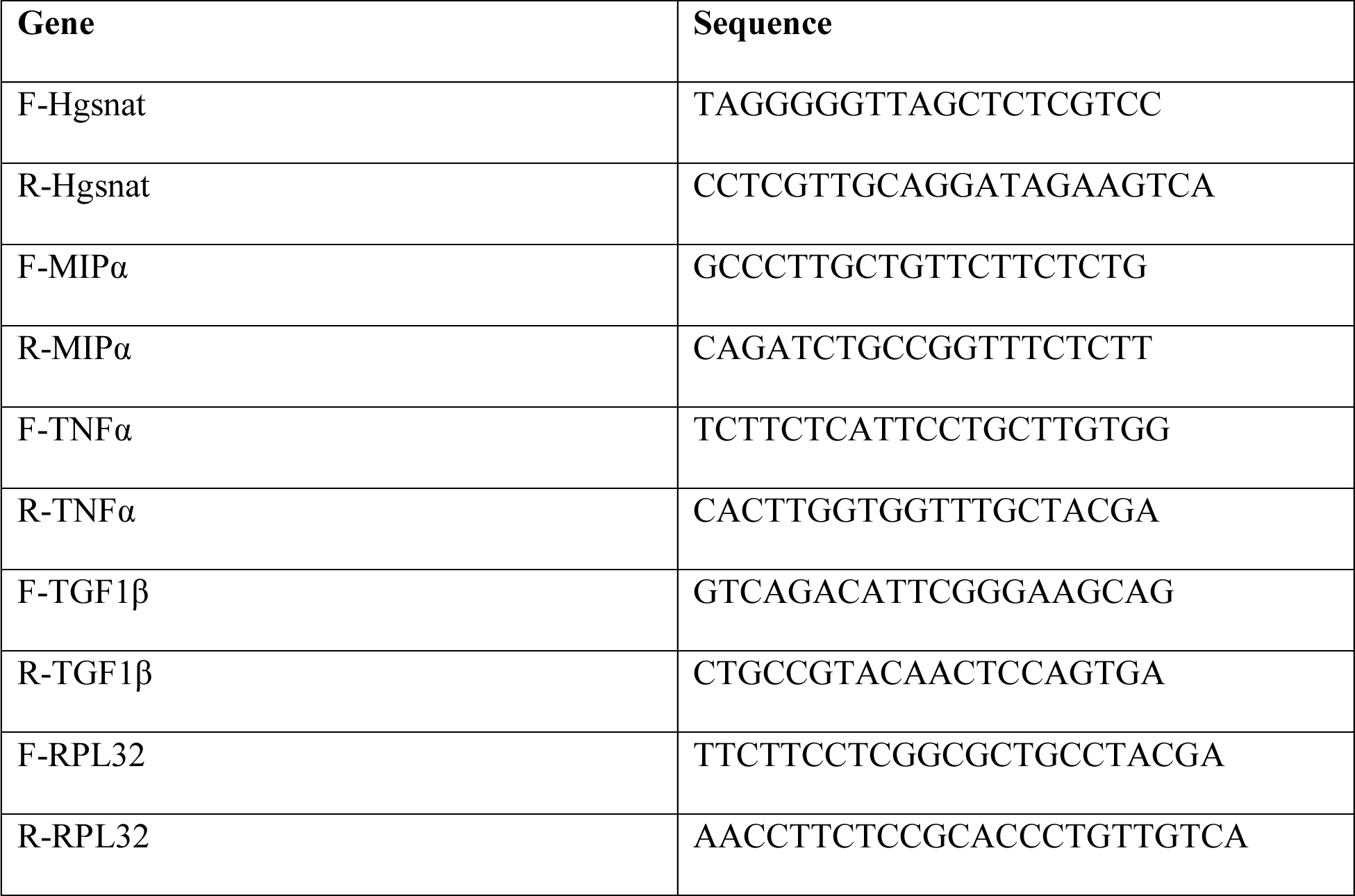

### RNA extraction and transcriptome sequencing

Total RNA was extracted from approximately 30 mg of hippocampal tissue using the RNeasy Mini Kit (Qiagen), according to the manufacturer’s instructions. RNA was quantified using the NanoDrop 8000. Samples with a 28S/18S ratio above 1.8, an OD 260/280 ratio greater than 1.9 and a RIN > 9 (Agilent Bioanalyzer 2100) were chosen for cDNA library construction. Library preparation and sequencing was made at the Institute for Research in Immunology and Cancer’s Genomics Platform (IRIC). All cDNA libraries were sequenced using single-end strategy (40 M reads per sample) on an Illumina NextSeq500 platform with a read length of 75 bp (single-end mode). Raw data were converted to FASTQ files using bcl2fastq (version 2.20) and allowed 0 mismatches in the multiplexing barcode. Data were analyzed by BioJupies using default parameters. Gene expression-level differences were accepted as statistically significant if they had a *p*-value of less than 0.05 and the result is visualized by an in-house MATLAB code.

### Normal Phase HPLC for glycosphingolipids in mouse brain extracts

Glycosphingolipids (GSLs) were analysed essentially as described previously [35]. Lipids from aqueous mouse brain homogenates (approximately 2.5 mg wet weight in 0.5 ml) were extracted with chloroform and methanol overnight at 4°C. The GSLs were then further purified using solid-phase C18 columns (Telos, Kinesis). After elution, the GSL fractions were dried down under a stream of nitrogen at 42°C and digested with recombinant endoglycoceramidase (rEGCase I, prepared by Genscript) to release the oligosaccharide headgroups. The liberated free glycans were then fluorescently-labelled with anthranillic acid (2AA). To remove excess 2AA label, labelled glycans were purified using DPA-6S SPE columns (Supelco). Purified 2AA-labelled oligosaccharides were separated and quantified by normal-phase high-performance liquid chromatography (NP-HPLC) as previously described [35]. The NP-HPLC system consisted of a Waters Alliance 2695 separations module and an in-line Waters 2475 multi λ-fluorescence detector set at Ex λ 360 nm and Em λ 425 nm. As a solid phase a 4.6 x 250 mm TSK gel-Amide 80 column (Tosoh Bioscience) was used. A 2AA-labelled glucose homopolymer ladder (Ludger) was included to determine the glucose unit values (GUs) for the HPLC peaks. Individual GSL species were identified by their GU values and quantified by comparison of integrated peak areas with a known amount of 2AA-labelled BioQuant chitotriose standard (Ludger). Protein concentration in homogenates was determined using the BCA assay (Sigma).

### Immunohistochemistry

Mouse brains were collected from animals perfused with 4% PFA in PBS and post-fixed in 4% PFA in PBS overnight. Brains were cryopreserved in 30% sucrose for 2 days at 4°C, embedded in Tissue-Tek® OCT Compound and stored at *−*80°C. Brains were cut in 40 µm-thick sections and stored in cryopreservation buffer (0.05 M sodium phosphate buffer pH 7.4, 15% sucrose, 40% ethylene glycol) at *−*20°C pending immunohistochemistry. Mouse brain sections were washed 3 times with PBS and permeabilized/blocked by incubating in 5% bovine serum albumin (BSA), 0.3% Triton X-100 in PBS for 1 h at room temperature. Incubation with primary antibodies, diluted in 1% BSA, 0.3% Triton X-100 in PBS, was performed overnight at 4°C. The antibodies used in the study and their working concentrations are shown in Table 1:

**Table 1:** Antibodies used in the study and their working concentration

| Antigen | Host/Target<br>species | Dilution | Manufacturer |
| --- | --- | --- | --- |
| Synapsin-1 | Rabbit anti-mouse | 1:200 | Abcam (ab64581) |
| GFAP | Rabbit anti-mouse | 1:300 | DSHB (8-1E7-s) |
| Lysosomal-associated<br>membrane protein 2<br>(LAMP2) | Rat anti-mouse | 1:200 | DSHB (ABL-93-s) |
| Heparan sulfate (10E4<br>epitope) | Mouse anti-mouse | 1:200 | AMSBIO (F58-10E4) |
| NeuN | Rabbit anti-mouse | 1:250 | Millipore Sigma (MABN140) |
| CD68 | Rabbit polyclonal to<br>CD68 | 1:200 | Abcam (ab125212) |
| G<sub>M2</sub> | Mouse humanized | 1:400 | KM966 |
| LC3B | Rabbit anti-mouse | 1:200 | GeneTex (GTX82986) |
| Vesicular glutamate<br>transporter 1 (VGLUT1) | Rabbit anti-mouse | 1:1000 (cells)<br>and 1:200<br>(tissues) | Abcam (ab104898) |
| Postsynaptic density<br>protein 95 (PSD-95) | Mouse anti-mouse | 1:1000 (cells)<br>and 1:200<br>(tissues) | Abcam (ab99009) |
| Ubiquitin | Mouse | 1:200 | Abcam (ab7254) |
| β-Amyloid (D54D2) | Rabbit anti-mouse | 1:200 | Cell Signaling (8243S) |
| Recombinant Anti-ATP synthase C antibody (SCMAS) | Rabbit anti-mouse | 1:200 | Abcam (ab181243) |
| Anti-O-GlcNAc | Mouse anti-mouse | 1:400 | Kerafast (EJH004) |
| Microtubule-associated protein 2 (MAP2) | Chicken anti-mouse | 1:2000 | Abcam (ab5392) |
| Vesicular gamma aminobutyric acid transporter vGAT | Rabbit anti-mouse | 1:1000 | Synaptic Systems (131003) |
| Gephyrin | mouse anti-mouse | 1:1000 | Synaptic Systems (147021) |
| Calreticulin | Rabbit anti-human | 1:200 | Sigma Aldrich (208910) |
| IgG | Goat anti-rabbit, anti-mouse, anti-rat or anti-chicken Alexa Fluor 488-, Alexa Fluor 555- or Alexa Fluor 633-conjugated | 1:400 | Thermo Fisher Scientific |

The mouse brain sections were washed 3 times with PBS and counterstained with Alexa Fluor-labeled secondary antibodies (dilution 1:400) for 2 h at room temperature. After washing 3 times with PBS, the mouse brain sections were treated with TrueBlack® Lipofuscin Autofluorescence Quencher (Biotium, 23007, dilution 1:10) for 1 min, and then again washed 3 times with PBS. The slides were mounted with Prolong Gold Antifade mounting reagent with DAPI (Invitrogen, P36935) and analyzed using Leica DM 5500 Q upright confocal microscope (10x, 40x, and 63x oil objective, N.A. 1.4). Images were processed and quantified using ImageJ 1.50i software (National Institutes of Health, Bethesda, MD, USA) in a blinded fashion. Panels were assembled with Adobe Photoshop.

### Immunocytochemistry

Cultured neurons were fixed in 4% paraformaldehyde and 4% sucrose solution in PBS, pH 7.4, for 20 min at day in vitro (DIV) 21. The cells were permeabilized with 0.1% Triton-X100 in PBS for 5 min, and non-specific binding sites were blocked with 5% BSA in PBS for 1 h and then, incubated overnight at 4°C with primary antibodies in 1% BSA in PBS (see Table 1 for the source of antibodies and their dilutions). On the following day, neurons were washed 3 times with PBS and labeled with Alexa Fluor 488- or Alexa Fluor-555- conjugated goat anti-rabbit or Alexa Fluor 633-anti-mouse IgG (1:1000, all from Thermo Fisher Scientific) for one hour at room temperature. Coverslips were washed 3 times again in PBS and mounted on slides using ProLong Gold mounting medium, containing 4′,6-diamidino-2-phenylindole (DAPI; Invitrogen, Cat # P36935), and analyzed by LSM510 Meta Laser or Leica TCS SPE confocal microscopes (× 63 glycerol immersion objectives, N.A. 1.4). Images were processed with Leica Application Suite X (LAS X) software and Photoshop 2021 (Adobe) and quantified using ImageJ 1.50i software (National Institutes of Health, Bethesda, MD, USA). Quantification was blinded and performed in at least 3 different experiments.

### Western blot

The cerebral cortical tissues were homogenized in five volumes of RIPA lysis buffer (50 mM Tris-HCl pH 7.4, 150 mM NaCl, 1% NP-40, 0.25% sodium deoxycholate, 0.1% SDS, 2 mM EDTA, 1 mM PMSF), containing protease and phosphate inhibitor cocktails (Sigma, cat# 4693132001 and 4906837001), using a Dounce homogenizer. The homogenates were kept on ice for 30 min and centrifuged at 13,000 g at 4°C for 25 min. The supernatant was centrifuged again at 13,000 g for 15 min, the protein concentration in resulting lysates was measured, and 20 µg of protein from each sample was separated by SDS-PAGE on 4–20% precast polyacrylamide gel (Bio-Rad, 4561096). Western blot analyses were performed according to standard protocols using the following antibodies: Anti-Grp78 (BiP) (rabbit polyclonal, 1:2000, Stressgen), CHOP (mouse, 1:50, DSHB), anti-ubiquitin, (1: 1000, rabbit, Sigma-Aldrich) and α-tubulin (1:2000, mouse, DSHB). Equal protein loading was confirmed by Ponceau S staining and normalized for tubulin immunoreactive band. Detected bands were quantified using ImageJ 1.50i software (National Institutes of Health, Bethesda, MD, USA).

### Production of the lentiviral vectors (LV) for expression of WT and mutant human HGSNAT

The plasmid expressing missense HGSNAT mutant P304L was obtained by site-directed mutagenesis of previously described pENTR1A-HGSNAT-GFP plasmid, containing the codon-optimized cDNA of WT human HGSNAT fused at the C-terminus with Green Fluorescent Protein (GFP) [36]. *HGSNAT-P311L-GFP* cDNA was further subcloned to lentiviral vector plasmid (pLenti CMV/TO Puro DEST (775-1), Addgene, Plasmid # 17432, ([37]), using Gateway^TM^ technology according to manufacturer’s instructions (Gateway™ LR Clonase™ II Enzyme mix, Invitrogen™ #11791020). In order to produce a lentiviral vector expressing HGSNAT-P311L-GFP protein under the control of CMV promotor, the HEK293T cells were co-transfected with pLenti-*HGSNAT-P304L*-*GFP* plasmid, the packaging plasmids REV (Addgene, Plasmid #12253) and pMDL (gag-pol) (Addgene, Plasmid #12251), and the envelope plasmid VSVG (Addgene, Plasmid#12259) in the presence of Lipofectamine 2000 Transfection Reagent (Invitrogen, 11668019). After 30 h, the cell medium containing lentivirus was collected and used to transfect primary mouse neurons, or human skin fibroblasts.

### Analysis of glycosaminoglycans by LC-MS/MS

Analysis of brain glycans was conducted as previously described by Viana *et al*. [38]. Briefly, 30- 50 mg of mouse brain tissues were homogenized in ice-cold acetone and centrifuged at 12,000× g for 30 min at 4°C. The pellets were dried, resuspended in 0.5 N NaOH and incubated for 2 h at 50°C. Then the pH of the samples was neutralized with 1 N HCl, and NaCl was added to the reaction mix in a final concentration of 3 M. After centrifugation at 10,000× g for 5 min at room temperature, the supernatants were collected and acidified using 1 N HCl. Following another centrifugation at 10,000× g for 5 min at room temperature, the supernatants were collected and neutralized with 1 N NaOH to a pH of 7.0. The samples were diluted at a ratio of 1:2 with 1.3% potassium acetate in absolute ethanol and centrifuged at 12,000× g and 4°C for 30 min. The pellets were washed with cold 80% ethanol, dried at room temperature, and dissolved in 50 mM Tris-HCl buffer. The samples were further filtered using AcroPrep^TM^ Advance 96-Well Filter Plates with Ultrafiltration Omega 10 K membrane filters (PALL Corporation, USA) and digested with chondroitinase B, heparitinase, and keratanase II, overnight at 37°C. The samples were analysed by mass spectrometry using a 6460 Triple Quad instrument (Agilent technologies) using Hypercarb columns, as described [38].

### Glucosamine treatment

Starting from 3 weeks of age, homozygous *Hgsnat^P304L^* male (n=24) and female (n=24) mice were randomly divided in two equal groups. The control group was administered water, while for the treatment group, water was supplemented with 10 mg/ml of glucosamine (Sigma, G4875-25G), which would result in a dose of approximately 2.0 g/kg BW/day. The dose formulation was changed twice a week. The untreated WT group included male and female siblings of *Hgsnat^P304L^* mice (n=14 and 13, respectively). At 16 weeks, all mice were studied by Y-maze and Novel Object Recognition behavioral tests and sacrificed to analyze CNS pathology as described above [39].

## Results

### 1. *Hgsnat^P304L^* mice show the complete deficiency of HGSNAT activity and storage of HS increased as compared with the *Hgsnat-Geo* knockout strain

The mouse *Hgsnat^P304L^* stain with the Pro304Leu mutation, an analog of human Pro311Leu variant [40] was generated as a murine model of MPS IIIC patients, with genetic changes leading to the synthesis of misfolded HGSNAT protein. The strain was produced essentially as described by Stephenson *et al.* [41], following the scheme shown in **Supplementary Figure S1**. Genotyping the offspring from heterozygous breeding revealed an expected Mendelian frequency (25%) for mice homozygous for the *Hgsnat^P304L^* allele. Similarly to previously described knockout *Hgsnat-Geo* mice [27], homozygous *Hgsnat^P304L^* mice of both sexes were viable, fertile, produced normal litter sizes, showed normal body weight gain and general behavior similar to their WT or heterozygous siblings until the age of 7-8 months, when they presented body weight loss, lethargy and signs of urinary retention.

HGSNAT protein could not be detected in neither WT nor *Hgsnat^P304L^* mouse tissues by Western blot using the commercially available antibodies because it is normally expressed at a very low level. Expression level of *Hgsnat* mRNA, measured by RT-q-PCR in the brain, liver, and kidney (**Figure 1A**) or by total RNA sequencing in the brain (not shown) of the homozygous *Hgsnat^P304L^* mice, was normal, and the message contained the expected c.911C>T change as demonstrated by Sanger sequencing of the RT-PCR products (**Figure 1A**). HGSNAT activity, measured with the fluorogenic substrate, 4-Muf-β-D-glucosaminide in the liver, kidney, and cultured embryonic skin fibroblasts (MEF cells) of homozygous *Hgsnat^P304L^* mice, was reduced to 0.3-5.0 % of that in WT mice, i.e. the levels similar to those measured in the tissues of homozygous knockout *Hgsnat-Geo* mice and below or close to the detection limit of the method (**Figure 1B**).

**Figure 1.**
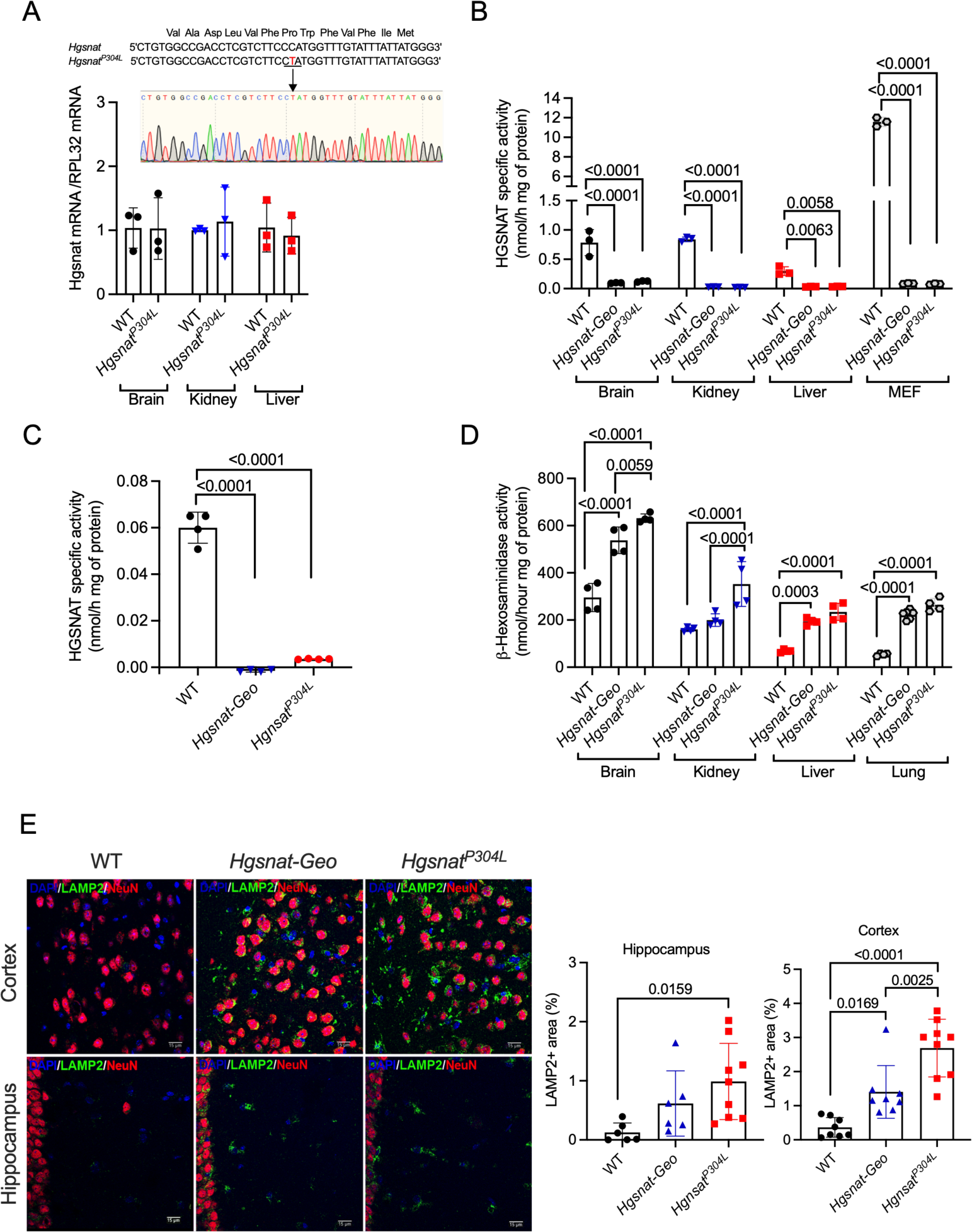
*Hgsnat^P304L^* homozygous mouse expresses normal levels of *Hgsnat* mRNA containing c. 911C>T mutation, shows complete deficiency of HGSNAT activity in tissues, and demonstrates bigger increase of lysosomal biogenesis in neurons as compared with the *Hgsnat-Geo* model. **(A)** Normal levels of *Hgsnat* mRNA containing c.911C>T mutation are expressed in the tissues of *Hgsnat^P304L^* mice. Total RNA was extracted from tissues of 4-month-old WT, *Hgsnat^P304L^* and *Hgsnat-Geo* mice and analyzed for *Hgsnat* expression by RT-qPCR. The values were normalized for the level of control *RPL32* mRNA. Data, means, and S.D. of experiments performed with 3 mice (2 male and 1 females) for each genotype are shown. All amplified PCR fragments were homozygous for c.911C>T mutation as verified by Sanger sequencing (inset). **(B)** HGSNAT specific activity was measured in the homogenates of brain, liver, and kidney of 4-months-old WT, homozygous *Hgsnat^P304L^* and *Hgsnat-Geo* mice, and in homogenates of cultured MEF cells, derived from homozygous animals of both strains using the fluorogenic substrate 4-Muf-β-D-glucosaminide. The activity of HGSNAT is reduced to the background level in all studied tissues of *Hgsnat^P304L^* and *Hgsnat-Geo* mice except for the brain. **(C)** HGSNAT specific activity was measured in the brain homogenates of 4-month-old WT, *Hgsnat^P304L^* and *Hgsnat-Geo* mice using BODYPI-glucosamine as a substrate. The activity of HGSNAT is reduced to the background level in the brains of both *Hgsnat^P304L^* and *Hgsnat-Geo* mice. **(D)** Specific activity of total lysosomal β-hexosaminidase in tissues of 4-months-old *Hgsnat^P304L^* and *Hgsnat-Geo* mice were measured, using the fluorogenic substrate 4-Muf-N-acetyl-β-D-glucosaminide. The activity of total lysosomal β-hexosaminidase shows a bigger increase in the tissues of *Hgsnat^P304L^* as compared with those of *Hgsnat-Geo* mice. **(E)** LAMP2 immunostaining is increased in neurons of 4-month- old *Hgsnat^P304L^* mice as compared with *Hgsnat-Geo* mice, suggesting higher levels of lysosomal storage. Lysosomes in neurons show a coarsely granular appearance. Panels show representative images of the somatosensory cortex (layers 4 and 5) and CA1 region of hippocampus of 4-month- old *Hgsnat^P304L^*, *Hgsnat-Geo* and WT mice. Bars represent 15 µm. Graphs show quantification of LAMP2 stained area by ImageJ software. All graphs show individual data, means and SD of experiments performed using tissues from 3 mice (or cell lines) per genotype. P values were determined by one-way ANOVA with Tukey’s multiple comparisons test (**A, C, E**) or two one-way ANOVA with Tukey’s multiple comparisons test (**B, D**).

In the brain tissues of both *Hgsnat^P304L^* and *Hgsnat-Geo* homozygous mice, the residual HGSNAT activity, measured against 4-Muf-β-D-glucosaminide, was reduced to 12-15% of normal. However, the N-acetyltransferase activity of the enzyme in the brain homogenates, directly measured using BODIPY-glucosamine [42], was reduced to below detection levels (**Figure 1C**). We, thus, conclude that similarly to human Pro311Leu HGSNAT [8], the mouse enzyme, containing the Pro304Leu variant lacks catalytic activity, while the residual activity against 4-Muf- β-D-glucosaminide in the brains of homozygous *Hgsnat^P304L^* and *Hgsnat-Geo* mice is caused by the action of an unknown hydrolase capable of cleaving non-acetylated β-D-glucosaminide. Interestingly, levels of total β-hexosaminidase activity, measured at 4 months in the brain, liver, kidney, and lungs of *Hgsnat^P304L^* mice, showed elevation compared with those in both WT and *Hgsnat-Geo* mice, consistent with increased levels of lysosomal biogenesis and lysosomal storage (**Figure 1D**). This was also supported by higher levels of Lysosome-Associated Membrane protein 2 (LAMP2)+ puncta in the somatosensory cortical (layers 4 and 5) and CA1 hippocampal neurons in the brains of 4-month-old *Hgsnat^P304L^* as compared with *Hgsnat-Geo* mice (**Figure 1E**).

To determine whether knock-in mice also show increased levels of GAG storage, we analyzed their brain tissues, urine, and dried blood spots by LC-MS/MS. This method measures the concentration of disaccharides derived from GAGs known to accumulate in MPS diseases: ΔDi-0S (dermatan sulfate), ΔDiHS-NS and ΔDiHS-0S (HS), as well as mono-sulfated keratan sulfate. Disaccharides were produced by specific enzyme digestion of each GAG and quantified by negative ion mode of multiple reaction monitoring (**Figure 2**). Analyses of the data have demonstrated drastically increased levels of HS-derived ΔDiHS-0S disaccharide in serum, urine, dry blood spots, and brain, and ΔDiHS-NS disaccharide in the dry blood spots and brain of homozygous *Hgsnat^P304L^* and *Hgsnat-Geo* mice. The levels of HS disaccharides in the dry blood spots and brain were significantly increased *Hgsnat^P304L^* mice as compared with *Hgsnat-Geo* mice, while in serum and urine they showed a trend for an increase. The levels of mono-sulfated KS were similar for *Hgsnat^P304L^* and *Hgsnat-Geo* mice and their WT counterparts, however, unexpectedly DS-derived ΔDi-0S showed a small but significant increase in brain tissues of *Hgsnat^P304L^* as compared with *Hgsnat-Geo* and WT mice of all ages.

**Figure 2.**
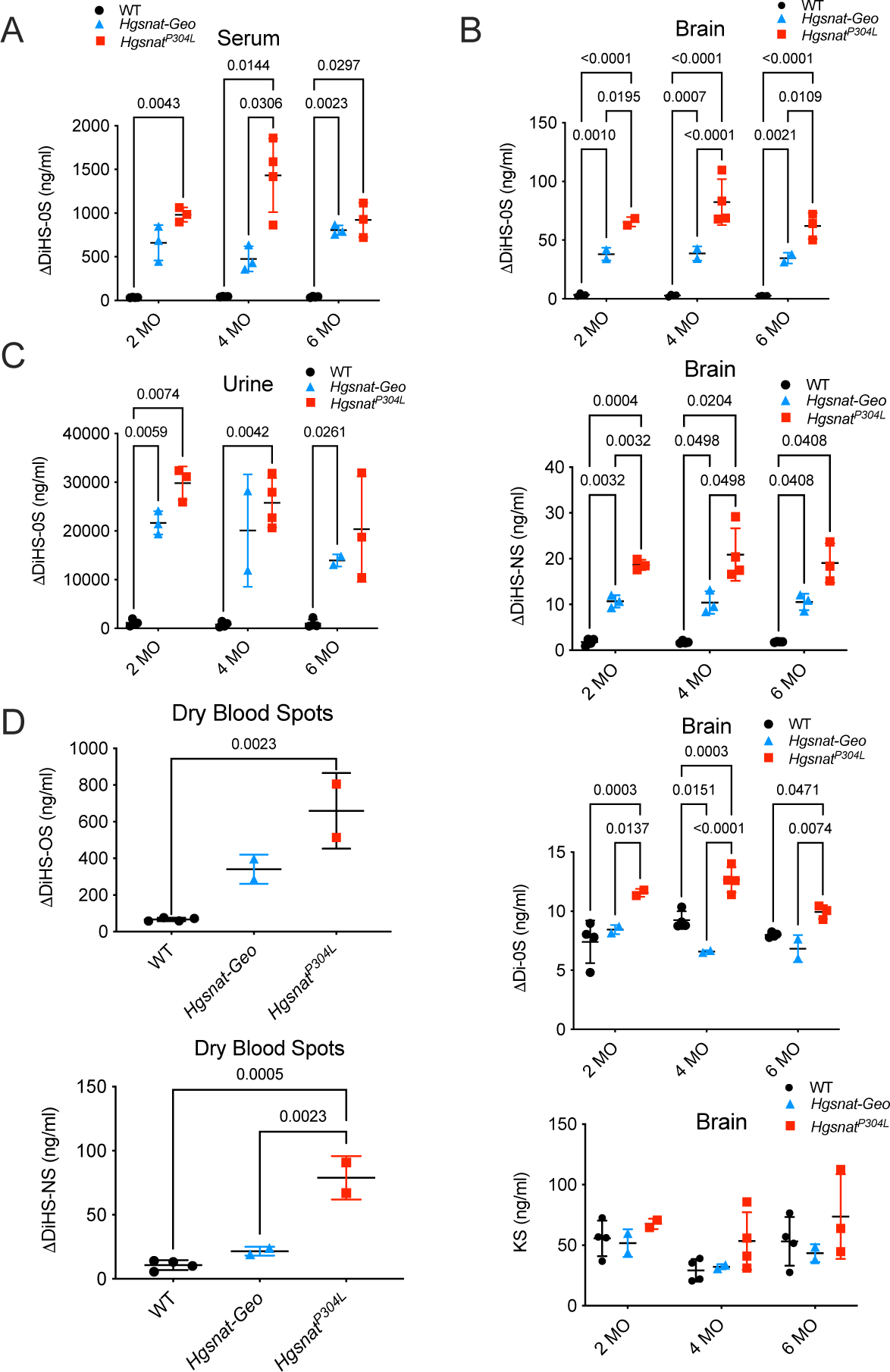
GAG levels in the tissues of MPS IIIC mice and controls. Levels of disaccharides produced by enzymatic digestion of DS (ΔDi-0S), HS (ΔDiHS-0S and ΔDiHS-NS), and mono-sulfated KS were measured by tandem mass spectrometry in blood serum. **(A)** brain tissues **(B)**, and urine **(C)** of WT, *Hgsnat^P304L^* and *Hgsnat-Geo* mice at the age of 2, 4 and 6 months as well as in the DBS **(D)** collected from WT, *Hgsnat^P304L^* and *Hgsnat-Geo* mice at the age of 2 months. All graphs show individual data, means, and SD of experiments performed using tissues from 3-4 mice per genotype. P values were measured using ANOVA with Tukey’s multiple comparisons test.

### 2. *Hgsnat^P304L^* mice show the earlier onset of behavioral changes, reduced longevity, and increased visceromegaly as compared with the *Hgsnat-Geo* strain

Previously, we have reported progressive behavioral changes in the homozygous *Hgsnat-Geo* mice, including hyperactivity and reduced anxiety revealed by the Open Field (OF) test at the age between 8 (6 in the female group) and 10 months, as well as deficits in spatial memory and learning capability, revealed by Morris Water Maze at the age of 10 months. To test whether increased lysosomal storage in *Hgsnat^P304L^* mice coincided with earlier onset of behavioral changes, mice were studied using the following behavioral tests: Elevated Plus Maze (EPM, anxiety, and fear), OF (anxiety and hyperactivity), Novel Object Recognition (NOR, short-term memory) and Y-maze (YM, short-term and spatial memory). The tests were performed every two months starting from the age of 2 months, each time with a naïve group of mice.

At the age of 4 months, both male and female *Hgsnat^P304L^* mice show abnormal behavior in all four tests, including increased hyperactivity (increased distance traveled in OF, **Figure 3A, B**), reduced anxiety (increased distance traveled in the central part of the arena in OF, **Figure 3A)**, increased percentage of time spent in the open arms and increased number of open arm entries in EPM, **Figure 3 C, D**), and deficits in spatial as well as in short-term memory (reduced recognition index in NOR, **Figure 3E**, reduced alteration rate in YM, **Figure 3F**). In contrast, *Hgsnat-Geo* mice demonstrate normal behavior in YM at both 4 and 6 months and show reduced alternation only at the age of 8 months (**Figure 3F**). Together, these data demonstrate that behavioral changes occur in *Hgsnat^P304L^* mice at least 2-4 months earlier than in the *Hgsnat-Geo* mice.

**Figure 3.**
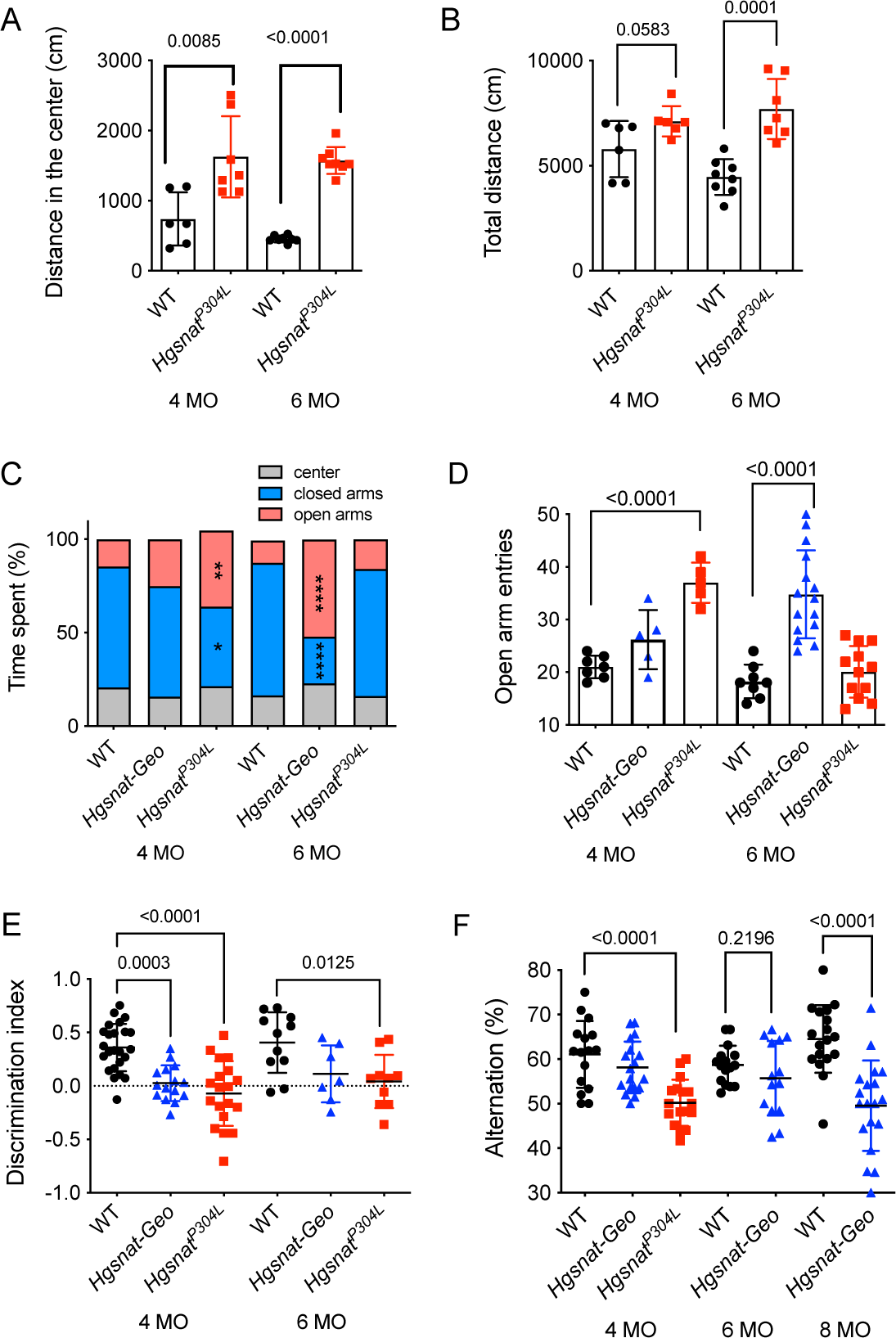
Neurobehavioral abnormalities in *Hgsnat-Geo* and *Hgsnat^P304L^* mice. Significant increase in the distance traveled in the central zone **(A)** and total distance traveled in the Open Field **(B)** by *Hgsnat^P304L^* mice as compared with age-matched WT controls. **(C)** Significant decrease and increase in the percent of time spent in open arms and closed arms in the Elevated Plus Maze, respectively, by *Hgsnat*-*Geo* and *Hgsnat^P304L^* mice as compared with age-matched WT controls. **(D)** Significant increase in the number of open arm entries in the Elevated Plus Maze by *Hgsnat*-*Geo* and *Hgsnat^P304L^* mice as compared to age matched WT controls. **(E)** Significant decrease in the discrimination index in *Hgsnat^P304L^* mice at 4 months and 6 months in Novel Object Recognition test as compared to age-matched WT controls. **(F)** *Hgsnat^P304L^* and *Hgsnat-Geo* mice show learning impairment in Y maze at 4 and 8 months, respectively. All graphs show individual data, means and SD of experiments performed with 5 or more mice per genotype. P-values were calculated by t-test for experiments involving comparison of 2 groups (**A** and **B**), and ANOVA with Tukey’s multiple comparisons test, when comparing 3 groups (**C-F**).

Similar to *Hgsnat-Geo* mice and the mouse models of MPS IIIA and MPS IIIB [43, 44], *Hgsnat^P304L^* mice develop urinary retention resulting in abdominal distension and requiring humane euthanasia. However, their average life span is ∼20 weeks less than the lifespan of *Hgsnat-Geo* mice (**Figure 4A**). Similar to *Hgsnat-Geo* mice, *Hgsnat^P304L^* animals do not develop skeletal abnormalities (**Supplementary Figure S3**). However, *Hgsnat^P304L^* mice sacrificed around the age of 40 weeks show enlargement of liver, kidneys, and spleen, unlike *Hgsnat-Geo* mice which show only hepatomegaly (**Figure 4B**).

**Figure 4.**
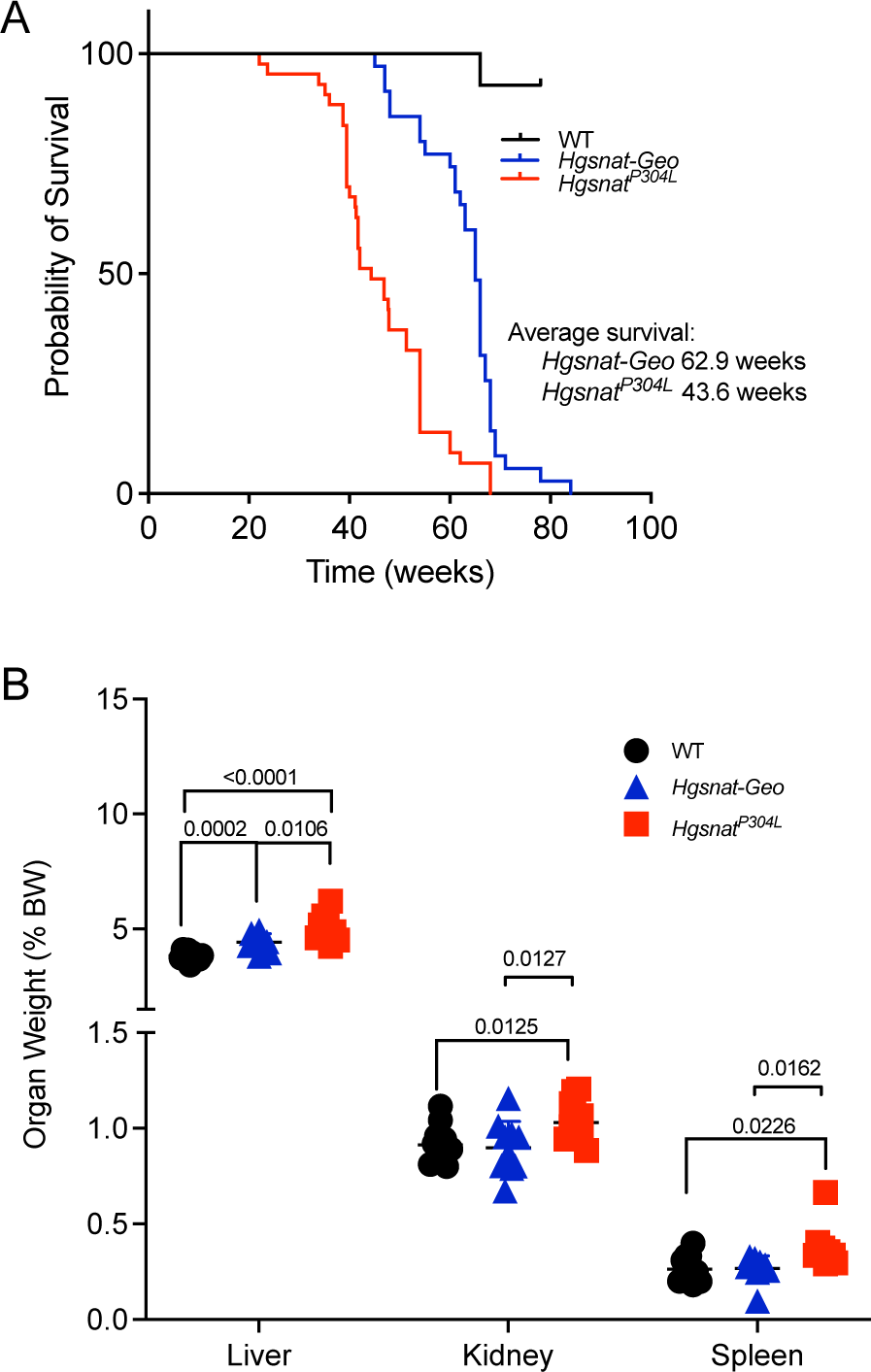
*Hgsnat^P304L^* mice have shorter life span than *Hgsnat-Geo* mice and show visceromegaly at the age of 8 months. **(A)** Kaplan-Meier plot showing survival of *Hgsnat^P304L^* (n=43) and *Hgsnat-Geo* male and female mice (*n*=35) and their WT counterparts (*n*=28). The significance of survival rate differences between strains was determined by the Mantel-Cox test (P<0.05). By the age of 45 weeks, most *Hgsnat^P304L^* mice had to be euthanized on the veterinarian request due to urinary retention, while *Hgsnat-Geo* mice survive to the average age of 63 weeks. **(B)** Wet organ weight of 8-months-old *Hgsnat^P304L^*, *Hgsnat-Geo* and WT mice is shown as a percentage of body weight. Enlargement of visceral organs, detected in *Hgsnat^P304L^* but not in *Hgsnat-Geo* mice as compared with age-matched WT controls, is consistent with the lysosomal storage and inflammatory cell infiltration previously reported for MPS III patients. All graphs show individual data, means and SD of experiments performed with 5 or more mice per genotype. P values were calculated using two-way ANOVA with post-hoc Tukey’s multiple comparison test.

### 3. *Hgsnat^P304L^* mice show more pronounced defects in synaptic neurotransmission when compared to the *Hgsnat-Geo* strain

To characterize synaptic neurotransmission in MPS IIIC mice, we performed whole-cell patch-clamp recordings on acute slices from *Hgsnat^P304L^* mice at P14-20 and P45-60. When the data were co-analysed together with our previous results for the age-matched groups of *Hgsnat-Geo* and WT mice [36], at both timepoints we found that the amplitudes of miniature excitatory postsynaptic currents mEPSC were significantly reduced in both *Hgsnat-Geo* and *Hgsnat^P304L^* mice as compared with WT mice. However, no significant difference was detected between *Hgsnat-Geo* and *Hgsnat^P304L^* mice (**Figure 5A and D**). Also, no differences in the kinetics of the mEPSCs between the two animal groups were observed (data not shown). Importantly, for both *Hgsnat-Geo* and *Hgsnat^P304L^* mice, there was an age-dependent (P14-20 vs P45-60) significant decrease in mEPSC amplitudes (**Figure 5A and D**) suggesting progressive synaptic deficits. The mEPSC frequency was significantly reduced in both *Hgsnat-Geo* and *Hgsnat^P304L^* mice as compared with WT controls at both ages, however, at P45-60, *Hgsnat^P304L^* mice displayed significantly reduced mEPSC frequencies as compared with *Hgsnat-Geo* mice (**Figure 5 B and C**). In contrast, aggravated defects in inhibitory neurotransmission were observed in the knock-in *Hgsnat^P304L^* mice as compared with the knockout model. At both P14-20 and P45-60, *Hgsnat^P304L^* mice showed significantly reduced frequencies of miniature inhibitory postsynaptic currents (mIPSC) as compared to both WT and *Hgsnat-Geo* mice of the same age (**Figure 5 E-H**).

**Figure 5.**
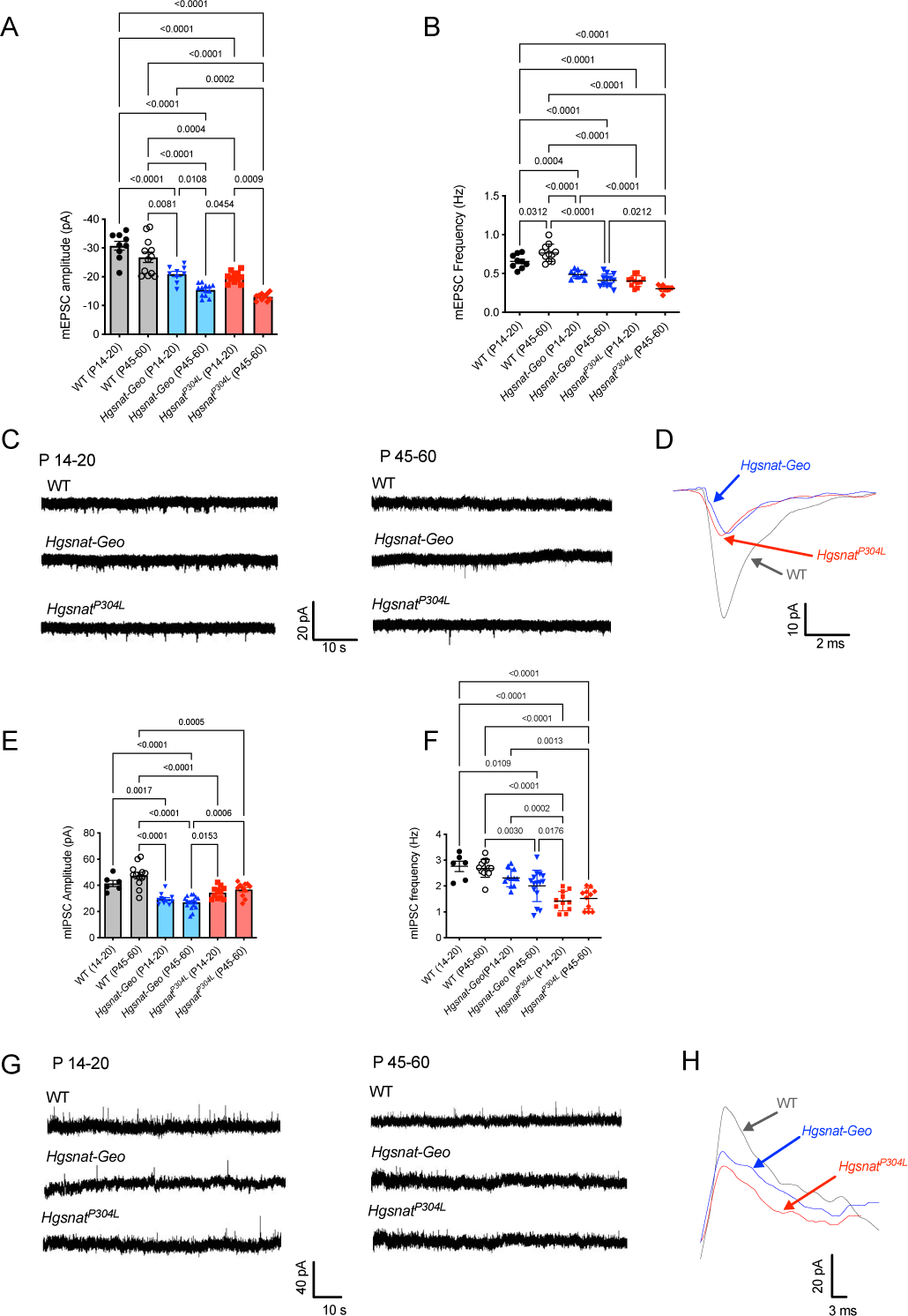
Synaptic neurotransmission is impaired in *Hgsnat-Geo* and *Hgsnat^P304L^* mice. Significant decrease in the amplitude(**A**) and frequency (**B**) of mEPSCs in *Hgsnat-Geo* and *Hgsnat^P304L^* mice at the ages of P14-20 and P45-60 as compared to age-matched WT controls. (**C**) Representative recordings of mEPSCs from WT, *Hgsnat-Geo* and *Hgsnat^P304L^* mice at P14-20 and P45-P60. (**D**) Overlay of representative individual mEPSC events from neurons of *Hgsnat-Geo*, *Hgsnat^P304L^* and WT mice. Significant decrease in the amplitude (**E**) and frequency (**F**) of mIPSCs in *Hgsnat-Geo* and *Hgsnat^P304L^* mice at the ages of P14-20 and P45-60 as compared with age-matched WT controls. (**G**) Representative recording of mIPSCs from neurons of WT, *Hgsnat-Geo,* and *Hgsnat^P304L^* mice at the ages of P14-20 and P45-60. (**H**) Overlay of representative individual mIPSC events from neurons of *Hgsnat-Geo*, *Hgsnat^P304L^* and WT mice. All graphs show individual data, means and SD of experiments performed with 6 or more mice per genotype. P values were calculated using one-way ANOVA with post-hoc Tukey’s multiple comparison test.

To test if changes in synaptic transmission were associated with those in the architecture of the synaptic compartment, we have analyzed hippocampal tissues of *Hgsnat-Geo* and *Hgsnat^P304L^* mice at the ages of 3 and 6 months by transmission electron microscopy (TEM). As earlier [36], we have measured the length and the area of post-synaptic densities (PSD) and the densities of synaptic vesicles in the terminals of symmetric (inhibitory) and asymmetric (excitatory) synapses of pyramidal CA1 neurons (**Figure 6A**).

**Figure 6.**
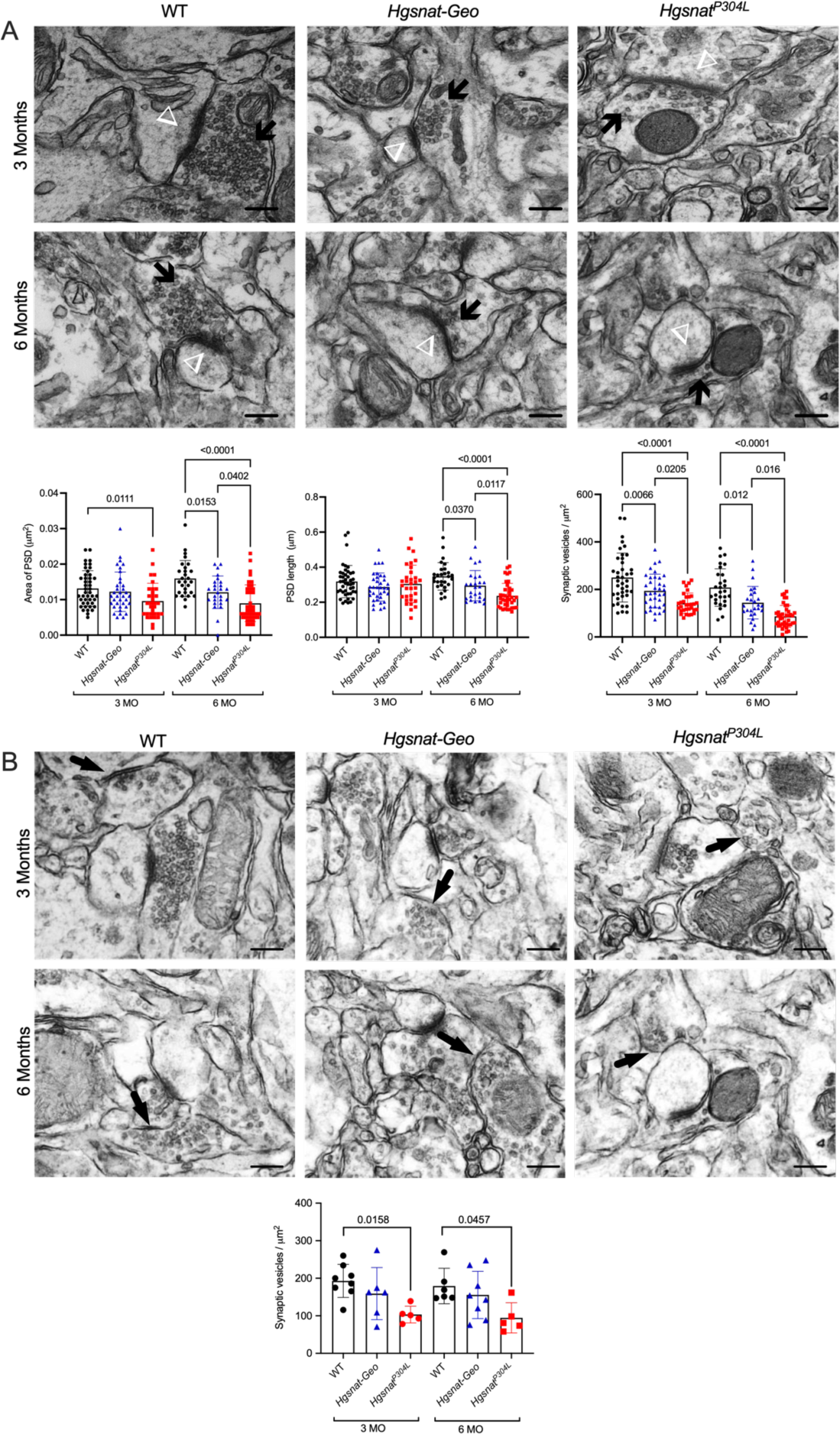
Reduction of synaptic vesicle densities, areas of postsynaptic densities, and length of postsynaptic densities in *Hgsnat-Geo* and *Hgsnat^P304L^* CA1 pyramidal hippocampal neurons. Quantification of density of synaptic vesicles, length (µm) and area of PSDs (µm^2^) in asymmetrical **(A)** and density of synaptic vesicles in symmetrical **(B)** pyramidal neurons from the CA1 region of the hippocampus. Synaptic terminals on the TEM images are marked with arrowheads. Data show values, means and SD of the results obtained with 3 different mice per genotype with 2-10 images quantified per animal. P-values were calculated by one-way ANOVA with Tukey post-hoc test for multiple comparisons. Scale bars equal 200 nm in all panels.

At the age of 3 months, the areas of excitatory PSDs of CA1 neurons in *Hgsnat-Geo* mice were similar to those in WT mice, while in the *Hgsnat^P304L^* mice they have already been significantly reduced. At the age of 6 months, the areas of excitatory PSDs in hippocampal neurons of both MPS IIIC mouse models were reduced as compared with WT mice, but the *Hgsnat^P304L^* mice expressed particularly drastic phenotype with PSD areas ∼50% smaller than those in WT mice. A similar trend was observed for the excitatory PSD length: by 6 months PSD length in *Hgsnat^P304L^* mice was significantly reduced as compared with both WT and *Hgsnat-Geo* mice. The density of synaptic vesicles (total number of synaptic vesicles in the axonal terminal divided by the area of the terminal in µm^2^) also showed a more rapid decrease in *Hgsnat^P304L^* mice with a reduction by ∼43% at 3 months and ∼60% at 6 months as compared with WT mice. In *Hgsnat-Geo* mice, they were reduced only by ∼32% and 41%, respectively (**Figure 6B**).

Together, these data revealed that the *Hgsnat^P304L^* strain shows more pronounced defects in synaptic (both excitatory and inhibitory) neurotransmission and synaptic architecture as compared with the *Hgsnat-Geo* strain.

### 4. *Hgsnat^P304L^* mice show accelerated progression in CNS pathology

Comparative analysis of pathological changes in the brain of *Hgsnat-Geo* and *Hgsnat^P304L^* mice demonstrated that they are aggravated in the knock-in mice. The levels of activated CD68+ microglia and GFAP+ astrocytes at the age of 4 months were significantly increased in the hippocampi and somatosensory (layers 4-5) cortices of *Hgsnat^P304L^* mice as compared with both WT and *Hgsnat-Geo* strains (**Figure 7A**). This coincided with the significantly increased expression levels of inflammatory cytokines MIP1α and TNFα in the brains of *Hgsnat^P304L^* as compared with *Hgsnat-Geo* mice (**Figure 7B**).

**Figure 7.**
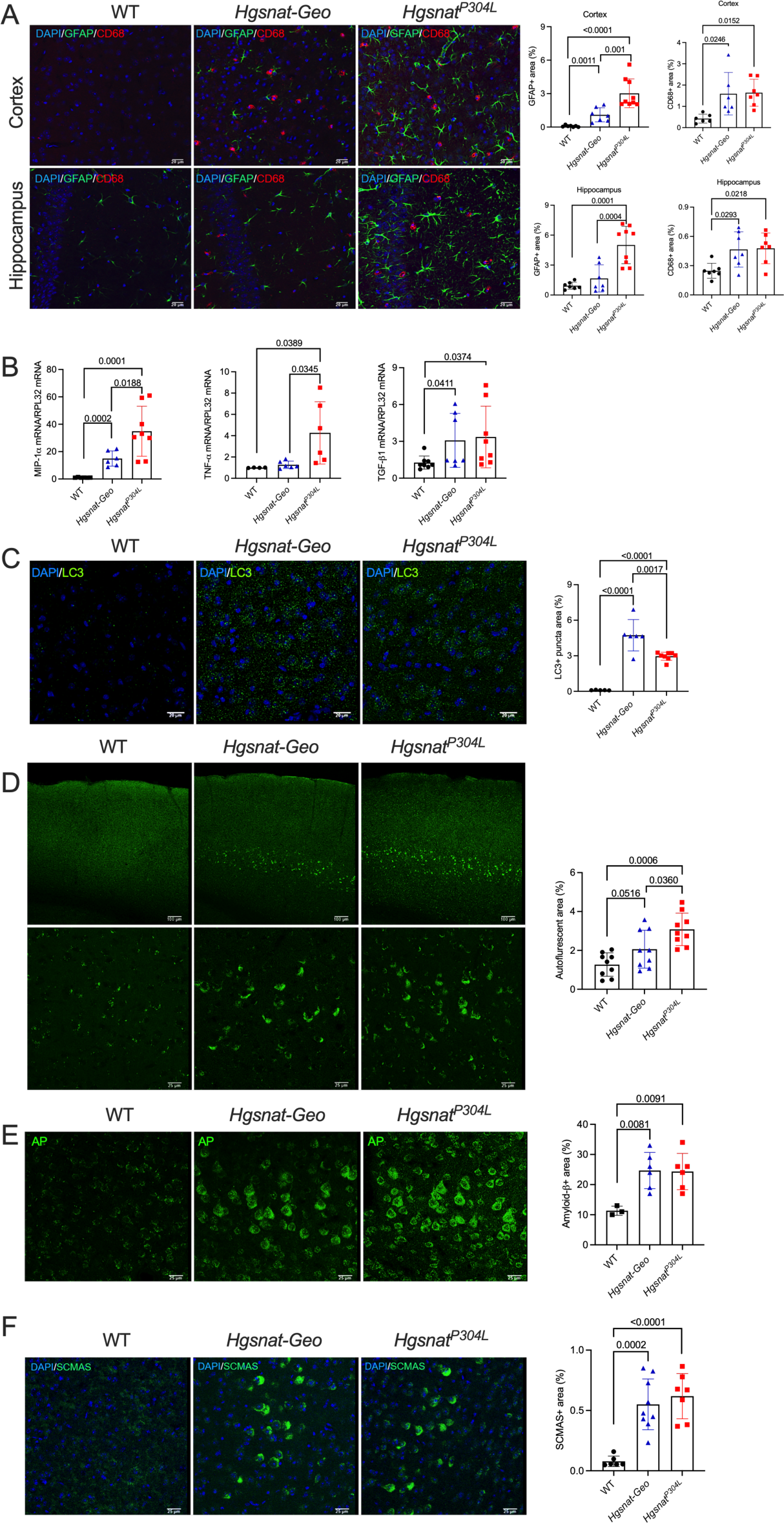
Aggravated pathological changes in the brains of *Hgsnat^P304L^* mice. **(A)** Astromicrogliosis in brain hippocampal and cortex regions of MPS IIIC mice is indicative of neuroinflammation. Panels show representative confocal microscopy images of brain tissues of 4-month-old *Hgsnat-Geo* and *Hgsnat^P304L^* mice and their age-matching WT controls stained with antibodies against CD68 (red) and GFAP (green) markers for activated microglia and astrocytes, respectively. DAPI (blue) was used as the nuclear counterstain. Activated microglia are marked with arrowheads and astrocytes, with asterisks; scale bar: 25 µm. Fluorescence was quantified with ImageJ software. Graphs show individual data, means, and SD from experiments performed with 3 mice per genotypes (3 areas/mouse). P values were quantified using one-way ANOVA with Tukey post-hoc test. **(B)** Total brain tissues of *Hgsnat^P304L^* mice show increased expression of inflammation markers, MIP1α (CCL3), and TNFα as compared with *Hgsnat-Geo* mice. Total RNA was isolated from whole mouse brain, reverse-transcribed to cDNA and quantified by RT- qPCR. The cytokine mRNA levels are normalized for the *RLP32* mRNA content. Data show individual data, means, and SD. At least three mice were analyzed for each age, sex and genotype. P values were calculated with one-way ANOVA with Tukey post-hoc test. Somatosensory cortices (layers 4-5) of *Hgsnat^P304L^* mice show increased levels of markers of impaired autophagy and proteolysis as compared with *Hgsnat-Geo* and/or age-matched WT mice: cytoplasmic LC3-positive puncta **(C),** granular autofluorescent ceroid materials **(D)**, amyloid protein (AP) **(E)**, and misfolded subunit C of mitochondrial ATP synthase (SCMAS) **(F)**. Panels show representative confocal microscopy images of brain tissues of 4-month-old (**A**, **B**, **D**) or 6-month-old (**C**, **E** and **F**) *Hgsnat-*Geo, *Hgsnat^P304L^* and WT mice. Bars represent 20 µm in **A** and **C**, 100 and 25 µm in **D,** and 25 µm in **E** and **F**. Graphs show results of quantification performed using ImageJ software. Individual data, means and SD obtained for 3-5 mice per genotypes (1-3 areas/mouse) are shown. P values were calculated using one-way ANOVA with Tukey post-hoc test.

The presence of LC3+ puncta was detected in layers 4-5 cortical pyramidal neurons of both *Hgsnat-Geo* and *Hgsnat^P304L^* mice but not of WT mice at the age of 6 months, suggesting an autophagy block (**Figure 7C**). Neurons of the same layers also contained increased levels of enlarged auto-fluorescent ceroid materials visible already at 4 months (**Figure 7D**). At 6 months neurons of the same layers, also, were heavily stained with the antibodies against the amyloid-*β* protein (AP) (**Figure 7E**) or misfolded subunit C of mitochondrial ATP synthase (SCMAS) (**Figure 7F**). Together, these data are suggestive of mitophagy block and a general impairment of proteolysis. Importantly, the number of cells containing autofluorescent material was significantly increased in *Hgsnat^P304L^* as compared with *Hgsnat-Geo* mice (**Figure 7D**), suggesting that the progression of this pathology is accelerated in the knock-in model.

To analyze if simple gangliosides GM2 and GM3, drastically increased in the brains of MPS IIIA-D patients [38] and in the knockout MPS IIIC mouse model [27], are also induced in the brain of *Hgsnat^P304L^* mice, glycosphingolipids were extracted from the pooled brain tissues of two, four, and six-months-old mice. Their glycan chains were fluorescently labelled with anthranilic acid and analyzed by normal-phase HPLC. Quantification of chromatograms (**Figure 8A**) demonstrated that brain glycosphingolipid composition was significantly altered in both *Hgsnat^P304L^* and *Hgsnat-Geo* mice but, on average, changes in the *Hgsnat^P304L^* mice were more pronounced. The most drastic changes were observed in the levels of GM3 (∼7-fold increase in *Hgsnat^P304L^* and ∼6-fold increase in *Hgsnat-Geo* mice), followed by GM2 (5- and 4-fold increase, respectively) and GA2 (3- and 2- fold increase, respectively). Interestingly, *Hgsnat^P304L^* mice showed a trend for a progressive increase in the levels of these gangliosides in contrast with *Hgsnat-Geo* mice where the levels remained similar at all studied ages. No changes were observed for complex gangliosides GM1a, GD1a, GD1b and GT1b.

**Figure 8.**
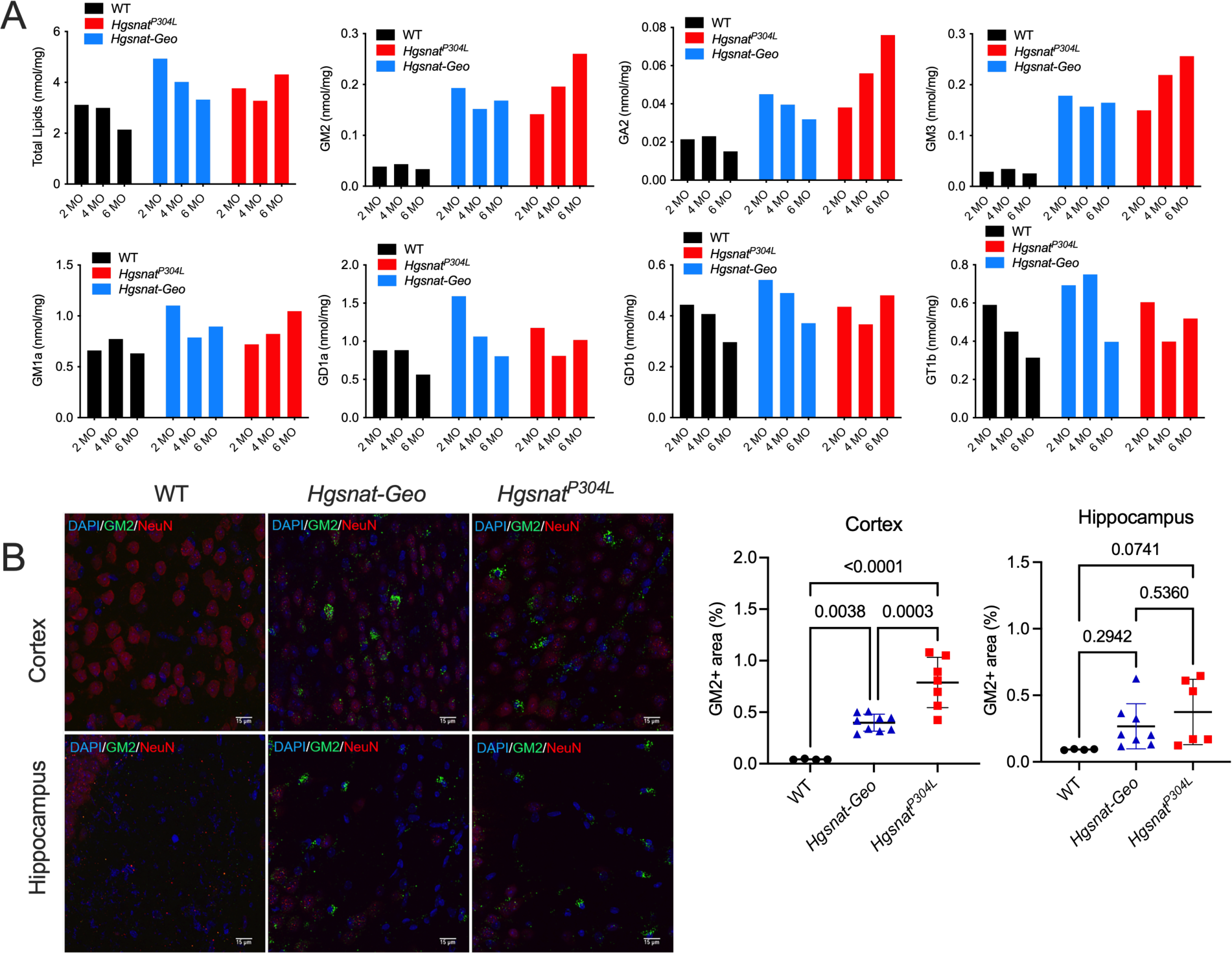
Alteration of sphingolipid levels in the brains of *Hgsnat-Geo* and *Hgsnat^P304L^* mice. **(A)** Levels of glycans produced by enzymatic cleavage of total sphingolipid extracts of brain tissues from *Hgsnat-Geo, Hgsnat^P304L^* and WT 2, 4 and 6-month-old mice were measured by normal HPLC. The values show % of the specific lipid. Pooled samples of 3 mice per age per genotype were analyzed. **(B)** Confocal microscopy images of brain cortex and hippocampus tissues of individual *Hgsnat- Geo, Hgsnat^P304L^* and WT 4-months-old mice stained with antibodies against GM2 (green) and NeuN (red). DAPI (blue) was used as the nuclear counterstain. Scale bar: 15 µm. Graphs show results of quantification performed using ImageJ software. Individual data, means, and SD obtained for 3 mice per genotypes (3 areas/mouse) are shown. P values were calculated using ANOVA with Tukey post-hoc test.

To confirm the HPLC results, we have analyzed the presence and distribution of GM2 ganglioside in brain tissues by immunohistochemistry, using the human-mouse chimeric monoclonal antibody, KM966 [45]. Numerous KM966+ neurons were present in the somatosensory cortex layers 4-5 and CA1 region of the hippocampus of both *Hgsnat^P304L^* and *Hgsnat-Geo* mice. However, in both brain regions the amounts of GM2+ cells were significantly increased in the knock-in as compared with knockout mice (**Figure 8B**).

### 5. Expression of the P304L HGSNAT variant in hippocampal cultured neurons of *Hgsnat-Geo* mice causes ER stress and aggravates deficits in the expression of synaptic proteins and synaptic architecture

To get insight into the molecular mechanisms underlying the severe phenotype of *Hgsnat^P304L^* mice, we have performed a bulk analysis of gene expression levels in the hippocampi of 4-months-old mice by RNA sequencing. Three mice (one female and two males) were analyzed for each genotype. The expression levels of each gene were compared between the *Hgsnat^P304L^* and *Hgsnat-Geo* strains, as well as between each of the MPS IIIC strains and the corresponding WT controls.

A higher number of hippocampal genes with altered expression levels was found in *Hgsnat^P304L^* (439 upregulated, 127 downregulated) as compared with *Hgsnat-Geo* mice (221 upregulated, 124 downregulated) (**Supplementary Figure S4**, **Table S1**). These genes were, then, classified according to their biological function and linked to metabolic or signaling pathways using automated GO (gene ontology terms) annotation [46]. The pathways involved in synaptic transmission (30-60% of all genes in the pathway), and neuronal growth and differentiation (10- 30% of all genes) showed major downregulation in both strains (**Figure 9A-B**). Importantly, the expression levels of the genes involved in GABAergic neurotransmission were reduced only in the *Hgsnat^P304L^* mice but not in *Hgsnat-Geo* mice, which was consistent with more pronounced defects in the inhibitory synapses detected in the knock-in mice by electrophysiology experiments (**Figure 9A**).

**Figure 9.**
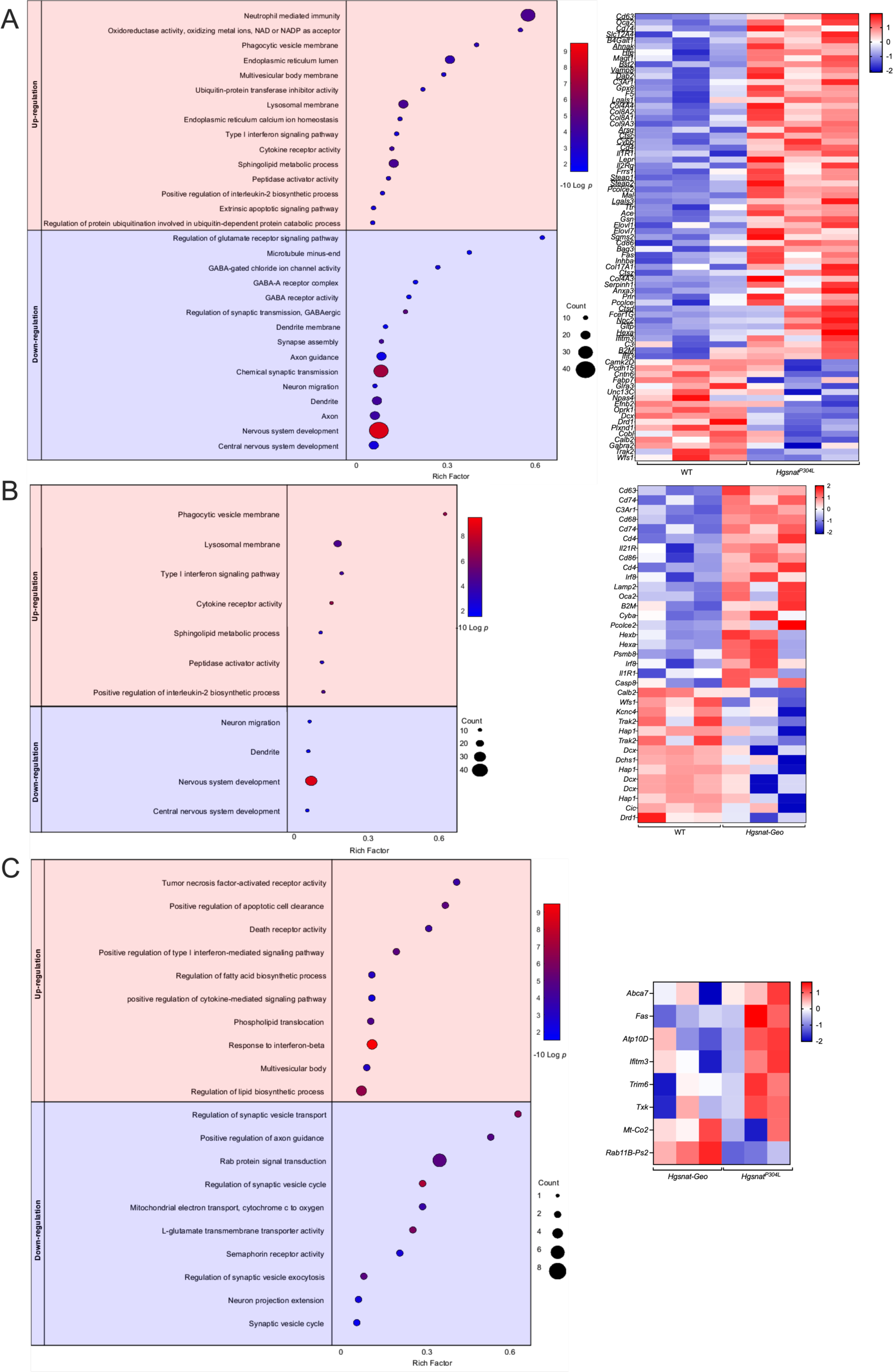
Hippocampal gene expression profiling of mRNAs in 4-month-old mice reveals increased expression of genes encoding lysosomal, lipid synthesis and pro-inflammatory proteins and reduced expression of genes involved in synaptic transmission, vesicular transport and neurogenesis in MPS IIIC mice. Dot plots (left) show significantly enriched GO terms (biological processes, molecular functions, and cellular components) and heat maps (right) of the genes significantly upregulated and downregulated in the hippocampi of *Hgsnat^P304L^* mice as compared with WT mice **(A)**, *Hgsnat-Geo* mice as compared with WT mice (**B**), and *Hgsnat^P304L^* mice as compared with *Hgsnat-Geo* mice (**C**). GO terms are plotted in the order of gene ratios, and each pathway is shown as a circle with the color representing the P-values (*-*log10) and the size representing the number of differentially expressed genes. The heatmap colors and their intensity show changes in gene expression levels. Data were obtained from sequencing of mRNA samples extracted from 3 different mice per genotype.

The most upregulated groups of genes were those encoding lysosomal and autophagosomal proteins, sphingolipid biosynthesis genes and genes involved in inflammatory and innate immune response, which was consistent with induced lysosomal biogenesis, alterations of ganglioside levels and inflammation observed in the mouse brains (**Figure 9A, B**). Specifically, several lysosomal (including *Arsg, Ctsc, Ctsz, Ctsd, Hexa, Npc2, Cd63, Cd74*, and *Slc12A4*) and inflammatory genes showed a significant increase only in *Hgsnat^P304L^* but not in *Hgsnat-Geo* mice mirroring higher levels of lysosomal storage and neuroinflammation in the 4-month-old knock-in animals (**Figure 9A**).

A direct comparison of *Hgsnat^P304L^* and *Hgsnat-Geo* expression profiles did not reveal significant changes in the expression of a gene or genes that could be directly responsible for the enhanced pathology in the knock-in mice. However, the pathways related to inflammation, cytokine production, apoptosis, and lipid biosynthesis were elevated, while those involved in synaptic function, neurogenesis, and mitochondrial biogenesis and function, downregulated in *Hgsnat^P304L^* as compared with *Hgsnat-Geo* mice. Besides, when compared with WT mice the levels of genes related to lysosomal/endosomal biogenesis (such as *Kcne2, Tfeb, Cst3, Gata2, Rilp, Pld1, Tlr7* and *Tmem59*) and inflammatory response (*Tlr7, Il-2, Il-25/Il-17, Ifngr1, Csf2rb2, Il15ra*, and *Il17rc*) showed a trend for a bigger increase in *Hgsnat^P304L^* than in *Hgsnat-Geo* mice (**Supplementary Figure S5)**. In similar fashion, genes involved in inhibitory synapse showed a trend for bigger reduction (**Supplementary Figure S5)**.

Interestingly a similar trend was observed for *Xbp1, Hspa5, Atf4, Ern1, Atf6, Atf3*, and *Hspa5* genes, which induction has been previously associated with the ER stress and unfolded protein response (UPR) (**Supplementary Figure S5**). This suggested a higher degree of the ER stress and UPR in the brain cells expressing the misfolded HGSNAT enzyme. To test this further, we have analyzed brain tissues by immunohistochemistry using antibodies against O-linked GlcNAc glycan and found increased levels of O-GlcNAc-modified proteins, an indication of the ER stress often associated with impaired cellular proteolysis [47], in the CA1 and cortical neurons of *Hgsnat^P304L^* as compared with *Hgsnat-Geo* mice (**Figure 10A**). We have also found increased levels of polyubiquitinated protein aggregates in the homogenates of dissected cortices of *Hgsnat^P304L^* as compared with *Hgsnat-Geo* mice (**Figure 10B**). This was consistent the increased number of pyramidal neurons containing ubiquitin-positive materials in somatosensory cortex layers 4 and 5 of *Hgsnat^P304L^* as compared with *Hgsnat-Geo* mice (**Figure 10C**). These results together confirmed higher levels of ER stress and UPR in the neurons of the knock-in MPS IIIC mice. At the same time, other markers of the ER stress, CHOP, and BiP, did not show an increase in the brains of *Hgsnat^P304L^* as compared with *Hgsnat-Geo* mice neither at 6 nor 8 months (**Supplementary Figure S6**).

**Figure 10.**
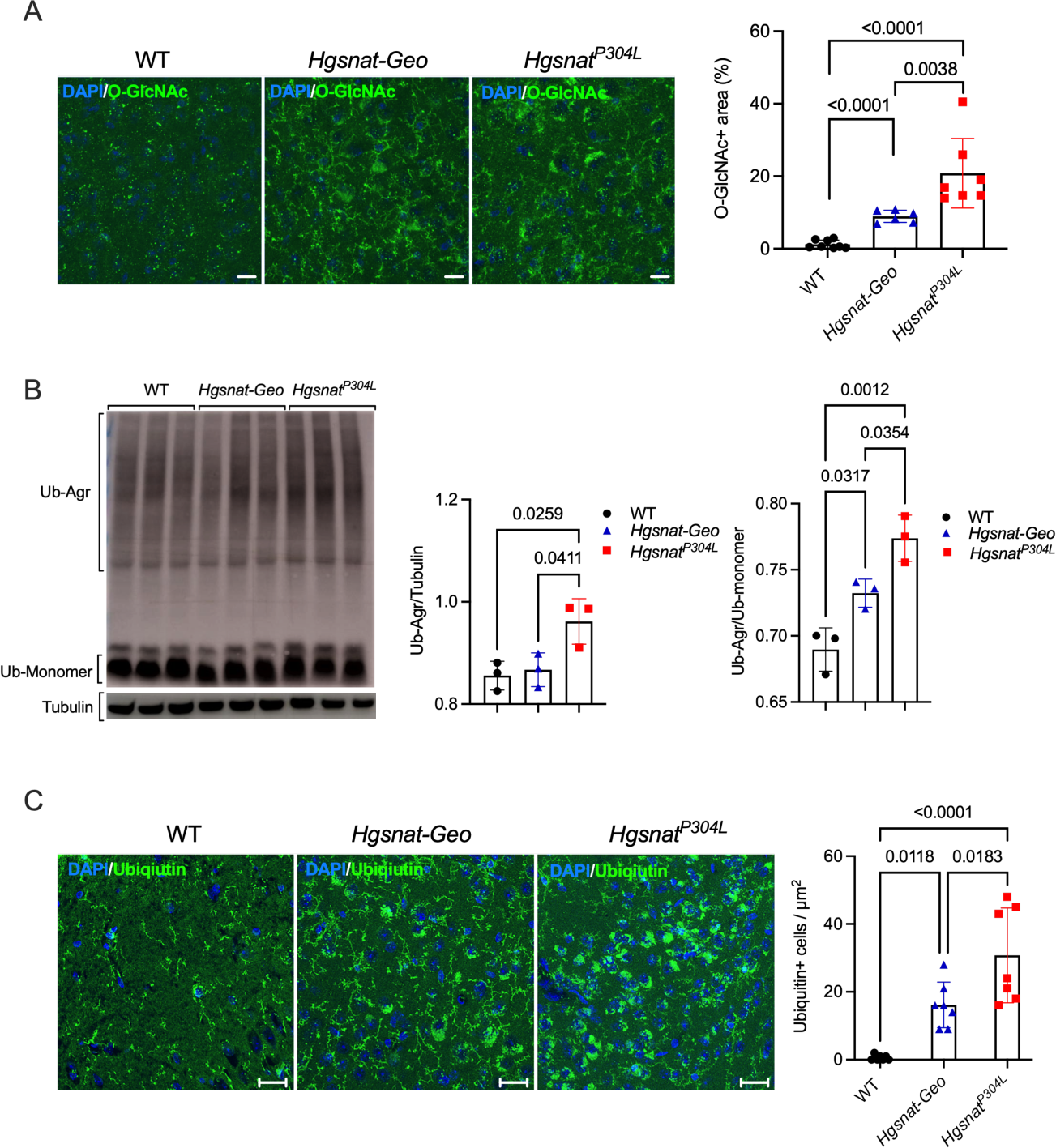
Levels of protein markers of unfolded protein response and ER stress are increased in the brains of *Hgsnat^P304L^* mice. **(A)** Brain cortex of *Hgsnat^P304L^* 6-month-old mice shows increased levels of O-GlcNAc-modified proteins as compared with *Hgsnat-Geo* and WT mice. Panels show representative images of brain cortex (layers 4-5) immunostained for O-GlcNAc (green). DAPI (blue) was used as a nuclear counterstain. Scale bar equals 25 µm. Three mice for each genotype were analyzed. Bar graph shows individual results, means and SD of quantification of O-GlcNAc staining with ImageJ software. Three mice per genotype (two areas per mouse) were analyzed. P values were calculated using ANOVA with Tukey post-hoc test. **(B)** Increased levels of ubiquitinated protein aggregates are detected in the brain homogenates of *Hgsnat^P304L^* mice by immunoblotting. Graphs show combined intensities (individual values, means, and SD) of protein ubiquitin+ bands quantified with ImageJ software and normalized by either intensity of tubulin bands or bands of ubiquitin monomers. Three mice per genotype were analysed. P values were calculated using ANOVA with Tukey post-hoc test. **(C)** Somatosensory cortex (layers 4-5) of *Hgsnat^P304L^* mice shows increased levels of pyramidal neurons containing cytoplasmic ubiquitin+ materials. Panels show representative confocal microscopy images of brain tissues of 6-month-old *Hgsnat-*Geo, *Hgsnat^P304L^* and WT mice stained for ubiquitin. Scale bar equals 25 µm. Graph shows results of quantification performed using ImageJ software. Individual data, means, and SD obtained for 4 mice per genotypes (1-3 areas/mouse) are shown. P values were calculated using one-way ANOVA test with Tukey post-hoc test.

To test directly whether the expression of the mutant Pro311Leu human HGSNAT variant aggravates neuronal dysfunction, we have expressed it in the primary cultured hippocampal neurons of *Hgsnat-Geo* mice. To confirm that the Pro311Leu variant indeed caused misfolding of the HGSNAT protein and its retention in the ER, we transduced HEK293 cells and human cultured skin fibroblasts with the LV vectors that encode the WT HGSNAT-GFP fusion protein and its Pro311Leu variant. Three days after transduction, cells expressing GFP marker were isolated by cell sorting and propagated. As expected, we detected highly increased HGSNAT activity in the cells overexpressing human WT HGSNAT, while the activity in the cells overexpressing the mutant variant was similar to that of non-transduced cells (**Supplementary Figure S7**). The WT protein was correctly processed in the lysosome as detected by the appearance of a 29-kDa band on the Western blot (**Supplementary Figure S7**). It was also showing a “halo-like” pattern around the LysoTracker Red stained lysosomes on the images obtained by a high-resolution confocal fluorescent microscopy suggesting that it was targeted to the lysosomal membrane (**Supplementary Figure S8**). In contrast, the mutant HGSNAT-GFP fusion protein was detected only in the form of a 75-kDa precursor (**Supplementary Figure S7**) and did not show any colocalization with LysoTracker Red or P115-stained Golgi apparatus. Instead, it was retained in the ER as demonstrated by its colocalization with the ER marker, Calreticulin (**Supplementary Figure S8**). In both fibroblasts (**Supplementary Figure S8A**) and HEK293 cells (data not shown), Pro311Leu HGSNAT-GFP protein was also forming cytoplasmic aggregates.

We further transduced primary hippocampal neurons of *Hgsnat-Geo* mice with either LV-HGSNAT-GFP or LV-P311L-HGSNAT-GFP to detect whether mutant protein expression aggravated defects in the expression of synaptic proteins or synaptic morphology. The cells were studied by immunocytochemistry to detect makers of synaptic vesicles (Syn1), GABAergic (VGAT/Gephyrin) and glutamatergic (VGLUT1/PSD-95) synapses. Non-transduced neurons of WT, *Hgsnat^P304L^*, and *Hgsnat-Geo* mice were studied for comparison.

Analysis of non-transduced cells confirmed that neurons from both *Hgsnat^P304L^* and *Hgsnat-Geo* mice showed a similar reduction in density of Syn1 puncta along the MAP2-stained dendrites as well as of PSD95-positive and VGLUT1-positive puncta (**Figure 11A-C**) as compared with WT neurons. This was consistent with our electrophysiology results demonstrating that glutamatergic synapse is similarly affected in both strains. In contrast, Gephyrin+/VGAT+ puncta were more reduced in neurons from *Hgsnat^P304L^* than in those from *Hgsnat-Geo* mice, confirming that the GABAergic synapse is further affected in the knock-in mice (**Figure 11D**).

**Figure 11.**
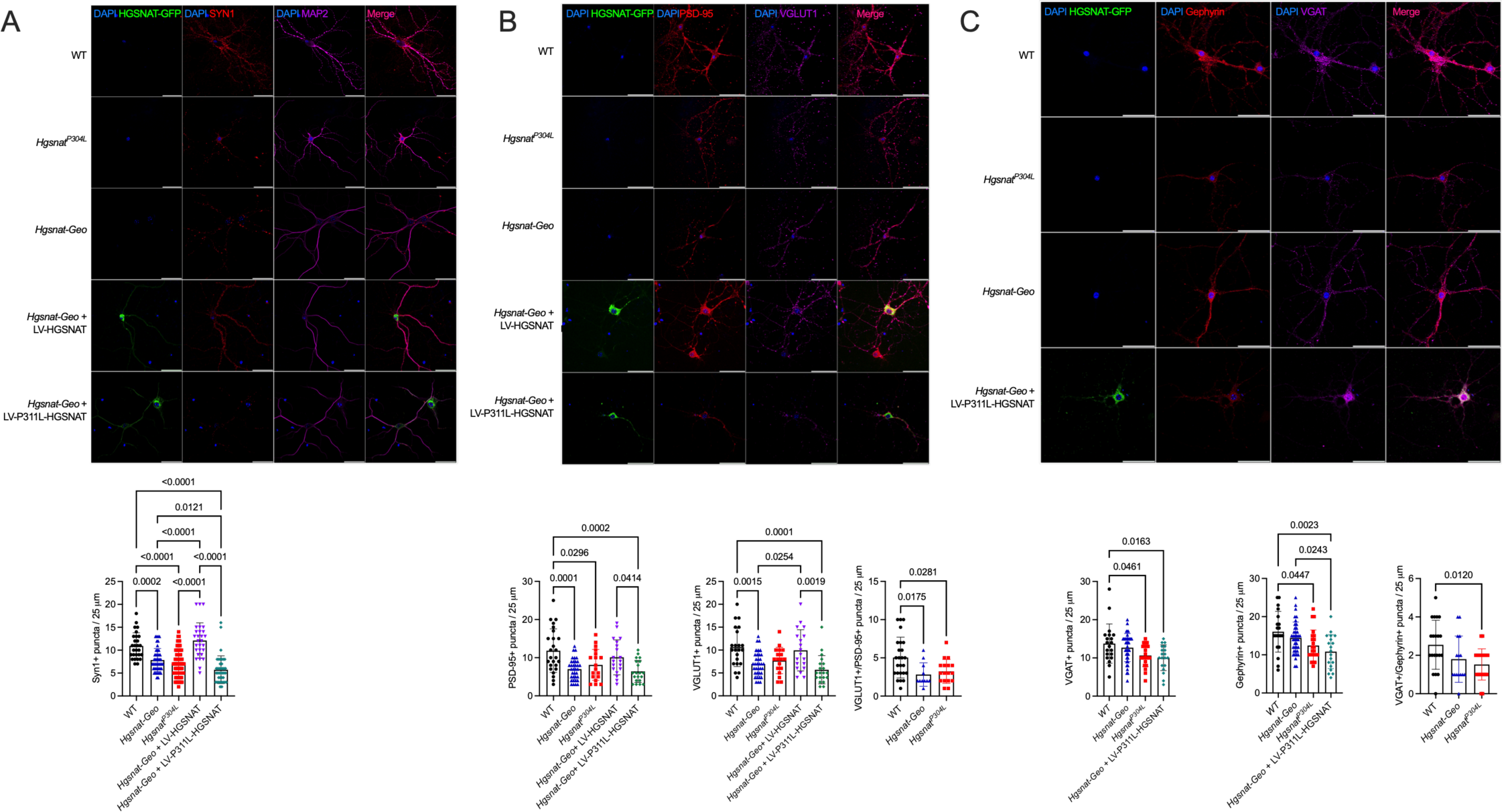
Expression of mouse P304L and human P311L mutant HGSNAT variants aggravates GABAergic synaptic defects in cultured primary hippocampal mouse neurons. **(A)** Hippocampal neurons from *Hgsnat^P304L^* and *Hgsnat-Geo* mice show equal reduction in density of Syn1+ puncta in proximity to MAP2+ dendrites as compared with WT cells. The levels of Syn1+ puncta are rescued by expression of WT active human HGSNAT but not of the P311L variant. **(B)** Hippocampal neurons from *Hgsnat^P304L^* and *Hgsnat-Geo* mice show an equal reduction in density of PSD-95+ and VGLUT1+ puncta and of PSD-95+/VGLUT1+ puncta in juxtaposition. The levels of PSD-95+ and VGLUT1+ puncta are rescued by expression of WT active human HGSNAT but not of the P311L variant. **(C)** Densities of Gephyrin+ and VGAT+ puncta and of Gephyrin+/VGAT+ puncta in juxtaposition are reduced in neurons from *Hgsnat^P304L^* but not from *Hgsnat-Geo* mice as compared with WT cells. Primary hippocampal neurons of *Hgsnat-Geo* mice transduced with LV-P311L-HGSNAT show reduction of Gephyrin+ and VGAT+ puncta as compared with WT and non-transduced *Hgsnat-Geo* cells. The panels show representative confocal images of neurons. Scale bars equal 50 µm. Graphs show results of puncta quantification with ImageJ software. Puncta were quantified in 20 µm-long segments of dendrite or axon, 30 µm away from the neuronal soma. Individual data, means and SD from 3 experiments, each involving pooled embryos from at least 3 mice per genotype are shown. P values were calculated using one-way ANOVA test with Tukey post-hoc test.

Transduction of neurons from *Hgsnat-Geo* mice with LV HGSNAT-GFP rescued levels of synaptic protein markers (dendrite-associated Syn1 puncta, VGLUT1+ puncta, and PSD-95+ puncta) (**Figure 11A-C**). In contrast, the *Hgsnat-Geo* neurons expressing mutant Pro311Leu HGSNAT-GFP protein showed levels of inhibitory (VGAT+, Gephyrin+) synaptic puncta significantly lower than those in the non-transduced cells (**Figure 11D**). The levels of excitatory (VGLUT+/PSD-95+) synapses in *Hgsnat-Geo* neurons expressing Pro311Leu HGSNAT-GFP showed a non-significant trend for reduction. Together, these results confirm that expression of Pro311Leu HGSNAT aggravates synaptic deficit caused by deficiency of HGSNAT activity and HS storage. Moreover, it expands the locus of the deficit toward the inhibitory GABAergic synapse.

### 6. Treatment of *Hgsnat^P304L^* mice with a pharmacological chaperon, glucosamine, partially restores the activity of the mutant enzyme and ameliorates clinical phenotype

Since we have shown that a competitive inhibitor of HGSNAT, glucosamine, rescues folding and activity of the missense HGSNAT variants in cultured skin fibroblasts from MPS IIIC patients[8], this drug was used here to test whether by reducing the load of the misfolded Pro304Leu protein we can ameliorate the clinical phenotype in *Hgsnat^P304L^* mice. Importantly, mice tolerate well chronic oral daily doses of glucosamine up to 2.0 g/kg BW and this compound can penetrate the brain parenchyma [48].

First, we tested if glucosamine exerts a chaperon effect in cultured MEF cells of *Hgsnat^P304L^* mice. The compound was added daily in a range of concentrations (3 to 10 Ki) to the cell culture medium, for a total of 5 days. Then, the cells were cultured overnight in the medium without the glucosamine, harvested, and lysed to measure the HGSNAT activity. Glucosamine in the concentration of 7 mM (10 Ki) had the maximal effect, increasing the residual activity about two-fold (**Supplementary Figure S9**). We further tested the drug in a group of 12 male and 12 female homozygous *Hgsnat^P304L^* mice. Starting from the age of 3 weeks (after weaning), mice were administered water supplemented with 10 mg/ml of glucosamine. Similar size control groups of WT and homozygous *Hgsnat^P304L^* mice were receiving normal drinking water. After 7 days, 3 mice from each group were sacrificed and HGSNAT activity measured in the homogenates of their brain tissues. The remaining animals were continued to be treated until the age of 16 weeks, when they were studied by the YM and NOR behavior tests and sacrificed at the age of 18 weeks to measure HGSNAT activity, HS levels and pathological changes in the CNS tissues. To test if chronic consumption of glucosamine resulted in major metabolic changes, the mouse body weight was measured monthly and blood glucose levels were analyzed before the sacrifice as described previously [49]. No difference between treated and untreated *Hgsnat^P304L^* mice was detected for both parameters (**Supplementary Figure S10A and B**), although, for an unknown reason, the blood glucose level in treated *Hgsnat^P304L^* mice was slightly lower than in untreated WT mice.

Brain residual levels of HGSNAT activity were significantly increased, although by a small margin, already after 7 days of treatment with glucosamine (**Figure 12A**). A similar increase was also observed in the brains of 5-months-old mice, after 13 weeks of treatment (**Figure 12B**). The level of HGSNAT activity measured in the liver of treated *Hgsnat^P304L^* mice was also increased as compared with that of untreated mice (**Figure 12C**). However, no difference in total *β*- hexosaminidase activity between treated and untreated mice was observed neither in the liver nor in the brain (**Figure 12D**).

**Figure 12.**
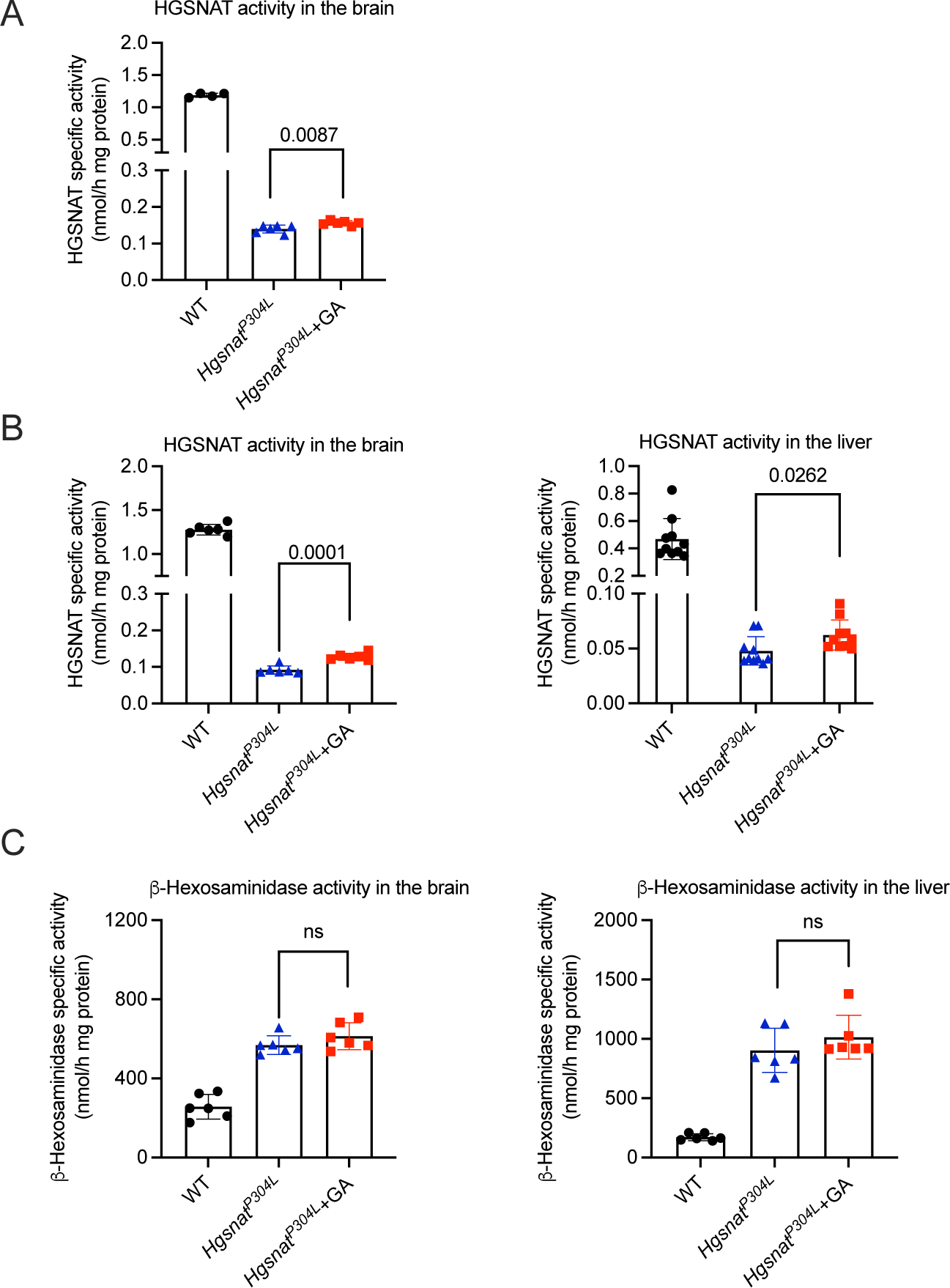
*Hgsnat^P304L^* mice treated with glucosamine show significant increase of HGSNAT activity in brain and liver tissues. HGSNAT activity, measured using the fluorogenic substrate, Muf-*β*-D-glucosaminide, was increased in the brain and liver tissue homogenates of 4-month-old *Hgsnat^P304L^* mice treated with glucosamine (+GA) for one **(A)** or 13 weeks **(B)** as compared with untreated *Hgsnat^P304L^* mice of the same age. No decrease of total *β*-hexosaminidase activity in both organs was detected in *Hgsnat^P304L^* mice treated with glucosamine for 13 weeks **(C)**. Individual results, means and SD from experiments performed with >3 mice per genotype, per treatment are shown. P values were calculated using an unpaired t-test.

When mouse memory and learning were tested using YM, we have observed a significant increase in the percent of alternation between the maze arms in treated mice, suggesting a rescue of the memory deficit. In the NOR test, WT mice spent more time exploring a novel than a familiar object showing a positive discrimination index, while untreated *Hgsnat^P304L^* mice spent equal time exploring both objects and, even, spent more time with a familiar object (negative discrimination index), showing signs of repetitive behavior (**Figure 13A**). For treated mice, we observed a significant increase in the discrimination index and the percent of time spent with a novel object (**Figure 13A**). Together, these data suggest that deficits in short-term memory in *Hgsnat^P304L^* mice were delayed by glucosamine treatment. Consistent with memory improvements we have also observed a partial rescue of the deficient protein markers of the excitatory synapse, PSD-95 and VGLUT1, in the hippocampal CA1 neurons and synaptic protein Syn1 in both hippocampal and cortical pyramidal neurons of treated *Hgsnat^P304L^* mice (**Figure 13B**).

**Figure 13.**
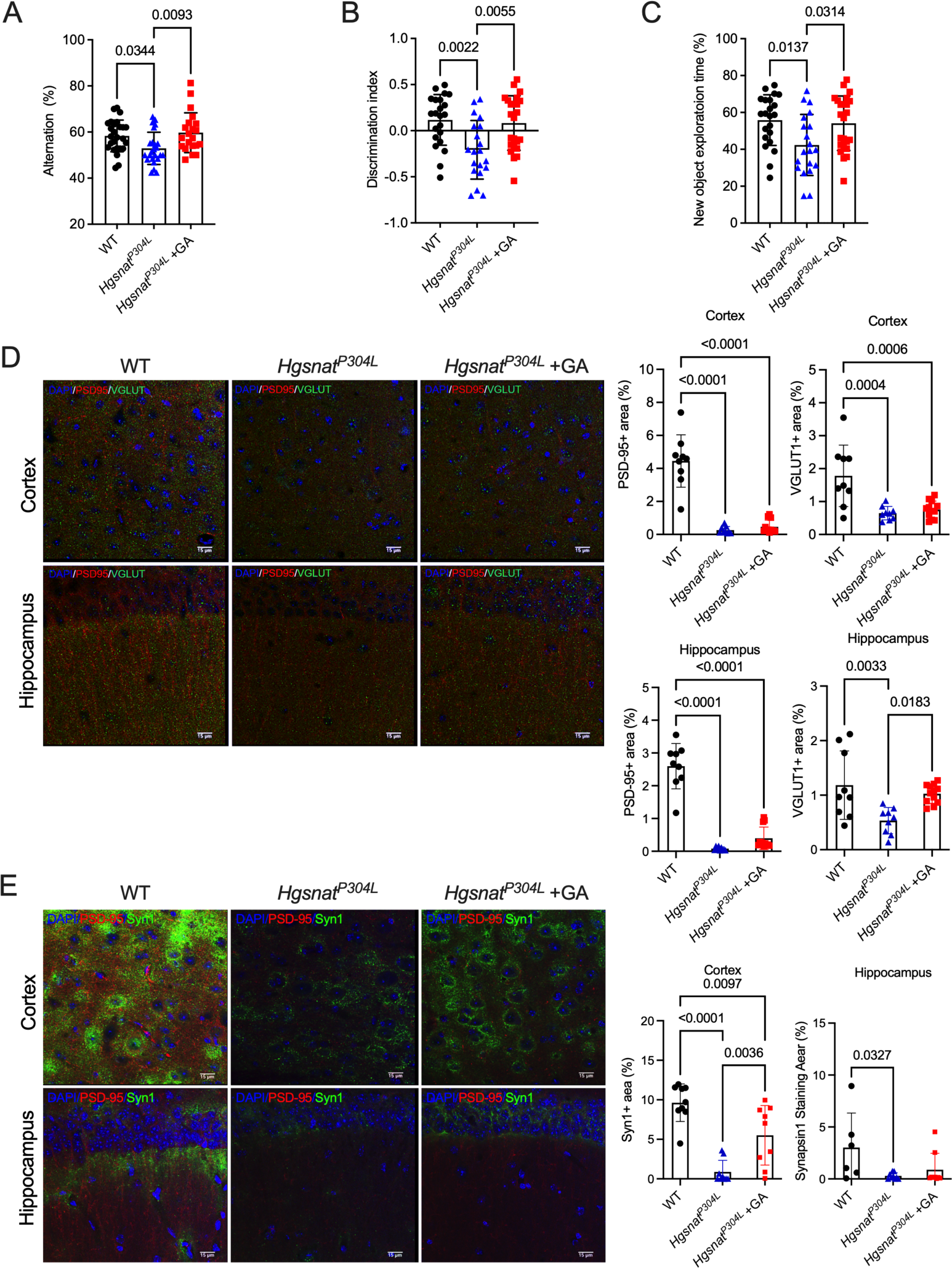
*Hgsnat^P304L^* mice treated with glucosamine reveal delay in development of deficits in memory and learning and partial rescue of synaptic protein markers in the CA1 area of the hippocampus. *Hgsnat^P304L^* mice, treated with glucosamine at the age of 4 months, show rescue or a trend for improvement of deficits in spatial/short-term memory in YM test **(A)** and short-term memory in NOR test **(B, C)** as compared with untreated *Hgsnat^P304L^* mice. Individual results, means and SD from experiments performed with 24 mice per genotype, per treatment are shown. P values were calculated using one-way ANOVA with Tukey post-hoc test. **(D)** Deficient levels of protein markers of glutamartergic synaptic neurotransmission, VGUT1 and PSD-95 are rescued in the CA1 hippocampal area of *Hgsnat^P304L^* mice, treated with glucosamine at the age of 4 months. Panels show representative images of brain cortex (layers 4-5) and CA1 area of the hippocampus of 4-month-old WT, and *Hgsnat^P304L^* mice treated or not with glucosamine stained for PSD-95 (red) and VGLUT1 (green). Scale bar equals 15 µm. The graph shows quantification of autofluorescence with ImageJ software. **(E)** Deficient level of synaptic vesicular protein Syn-1 are rescued in the somatosensory cortex area of *Hgsnat^P304L^* mice, treated with glucosamine at the age of 4 months. Panels show representative images of brain cortex (layers 4-5) and CA1 area of the hippocampus of 4-month- old WT, and *Hgsnat^P304L^* mice treated or not with glucosamine stained for PSD-95 (red) and Syn1 (green). Scale bar equals 15 µm. The graph shows quantification of autofluorescence with ImageJ software.

To test if delayed memory impairment in treated *Hgsnat^P304L^* mice coincided with a reduction in the development of brain pathology, we have analyzed fixed brain slices of treated and untreated *Hgsnat^P304L^* mice, as well as their WT counterparts, for the markers of primary and secondary storage (HS/LAMP2 and GM2 ganglioside, respectively), micro- and astrogliosis (CD68 and GFAP, respectively), and misfolded proteins accumulation (SCMAS, auto-fluorescent ceroid materials, and O-GlcNAc-modified proteins). These biomarkers were prioritized because, as described above, they were a key for discriminating between the aggravated phenotype of *Hgsnat^P304L^* and a milder phenotype of *Hgsnat-Geo* mice. Besides, we studied the expression levels of inflammatory cytokines, TNF-*α*, and MIP-1*α*, GAG levels, and the levels of ubiquitinated protein aggregates.

We found that some pathological signs were rescued or partially rescued in the treated mice. The pathological changes showing the best response to glucosamine treatment were an accumulation of autofluorescent ceroid materials and misfolded SCMAS in the deep cortical pyramidal neurons (**Figure 14A and B**), and accumulation of GM2 ganglioside in pyramidal neurons of somatosensory cortex and CA3 area of the hippocampus (**Figure 14C**). All three biomarkers were reduced in treated mice almost to the levels observed in the WT mice of the similar age. The level of total brain HS measured by the LC-MS/MS analysis showed a general trend for reduction in the mouse brain homogenates, however still remained at the significantly higher level than that in the WT mice (**Figure 15A**). On the other hand, immunohistochemistry has revealed a significant reduction of HS+ areas in the pyramidal neurons of both cortex and hippocampus (**Figure 15B**). The size and abundance of lysosomes (LAMP2+ area) showed significant reduction only in the cortex and just a trend for reduction in the hippocampus (**Figure 15B**). At the same time, glucosamine treatment did not change levels of astromicrogliosis, one of the hallmarks of CNS pathology in MPS IIIC and other neurological MPS diseases. The levels of CD68+ cells were similar for untreated and treated mice in the cortex and hippocampus, whereas GFAP+ cells were unchanged in the cortex, but, for an unknown reason, increased in the hippocampus of treated mice (**Supplementary Figures S11**). The levels of ubiquitinated protein aggregates, and O-GlcNAc-modified proteins were undetectable in the brains of both treated and untreated *Hgsnat^P304L^* mice as well as of WT mice at 4 months of age (data not shown).

**Figure 14.**
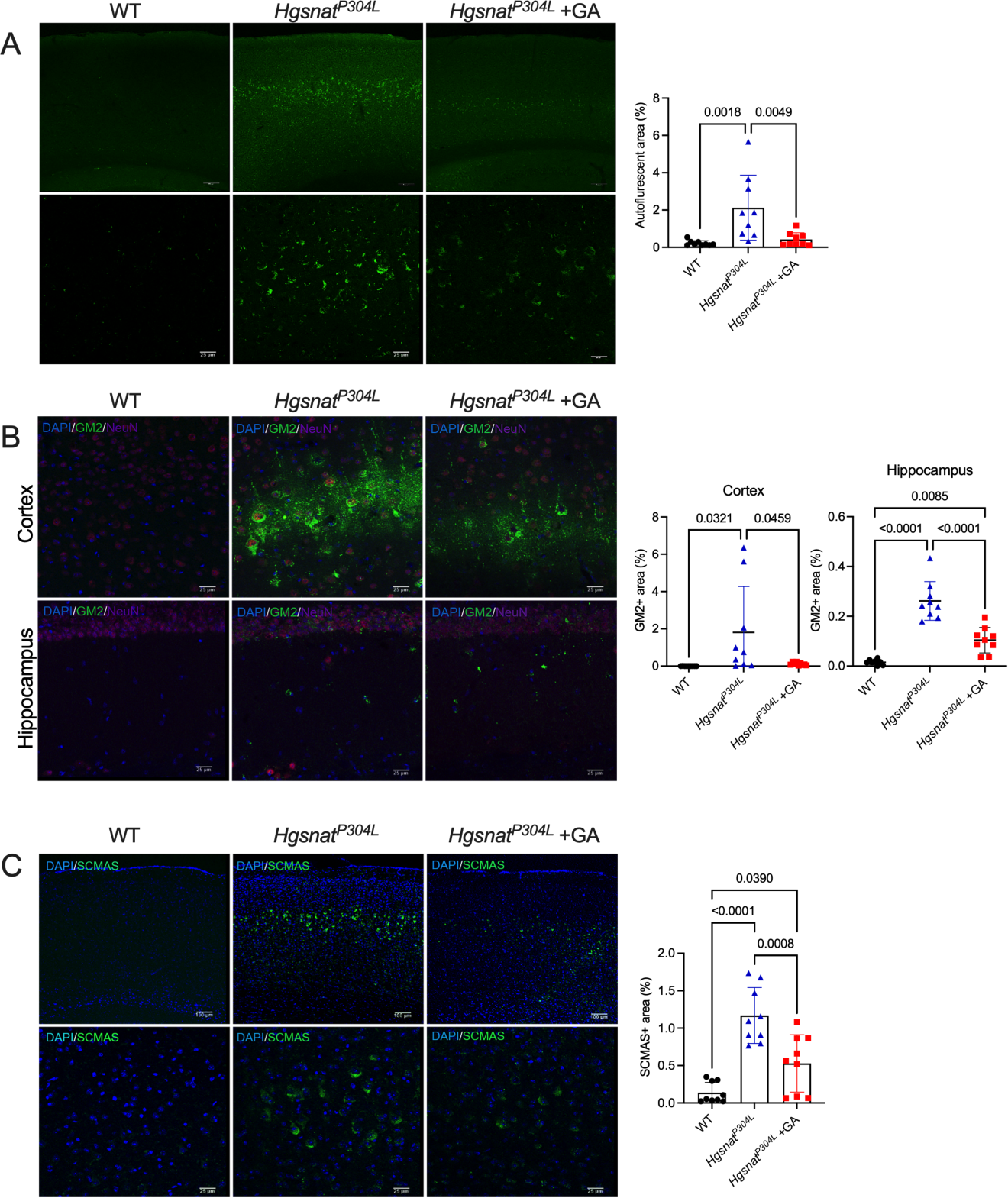
Biomarkers of CNS pathology are improved in the cortex of *Hgsnat^P304L^* mice treated with glucosamine. **(A)** Reduction of granular autofluorescent ceroid in cortical neurons. Panels show representative images of brain cortex (layers 4-5) of 4-month-old WT, and *Hgsnat^P304L^* mice treated or not with glucosamine showing autofluorescent ceroid inclusions in the neurons (green). Scale bar equals 100 µm (upper panels) and 25 µm (lower panel). The graph shows quantification of autofluorescence with ImageJ software. **(B)** Reduction of GM2 ganglioside storage in cortical neurons. Panels show representative images of brain cortex (layers 4-5) of 4-month-old WT, and *Hgsnat^P304L^* mice treated or not with glucosamine showing immunostaining for GM2 ganglioside and NeuN. DAPI was used as a nuclear counterstain. Scale bar equals 100 µm (upper panels) and 25 µm (lower panel). The bar graph shows quantification of GM2 staining with ImageJ software. **(C)** Reduction of misfolded SCMAS (subunit C of mitochondrial ATP synthase) aggregates in cortical neurons. Panels show representative images of brain cortex (layers 4-5) of 4-month-old WT and *Hgsnat^P304L^* mice treated or not with glucosamine immunostained for SCMAS. DAPI (blue) was used as a nuclear counterstain. Scale bar equals 100 µm (upper panels) and 25 µm (lower panel). The bar graph shows quantification of SCMAS staining with ImageJ software. All graphs show individual data, means and SD obtained for 3 mice per genotypes (3 areas/mouse). P values were calculated using one-way ANOVA test with Tukey post-hoc test.

**Figure 15.**
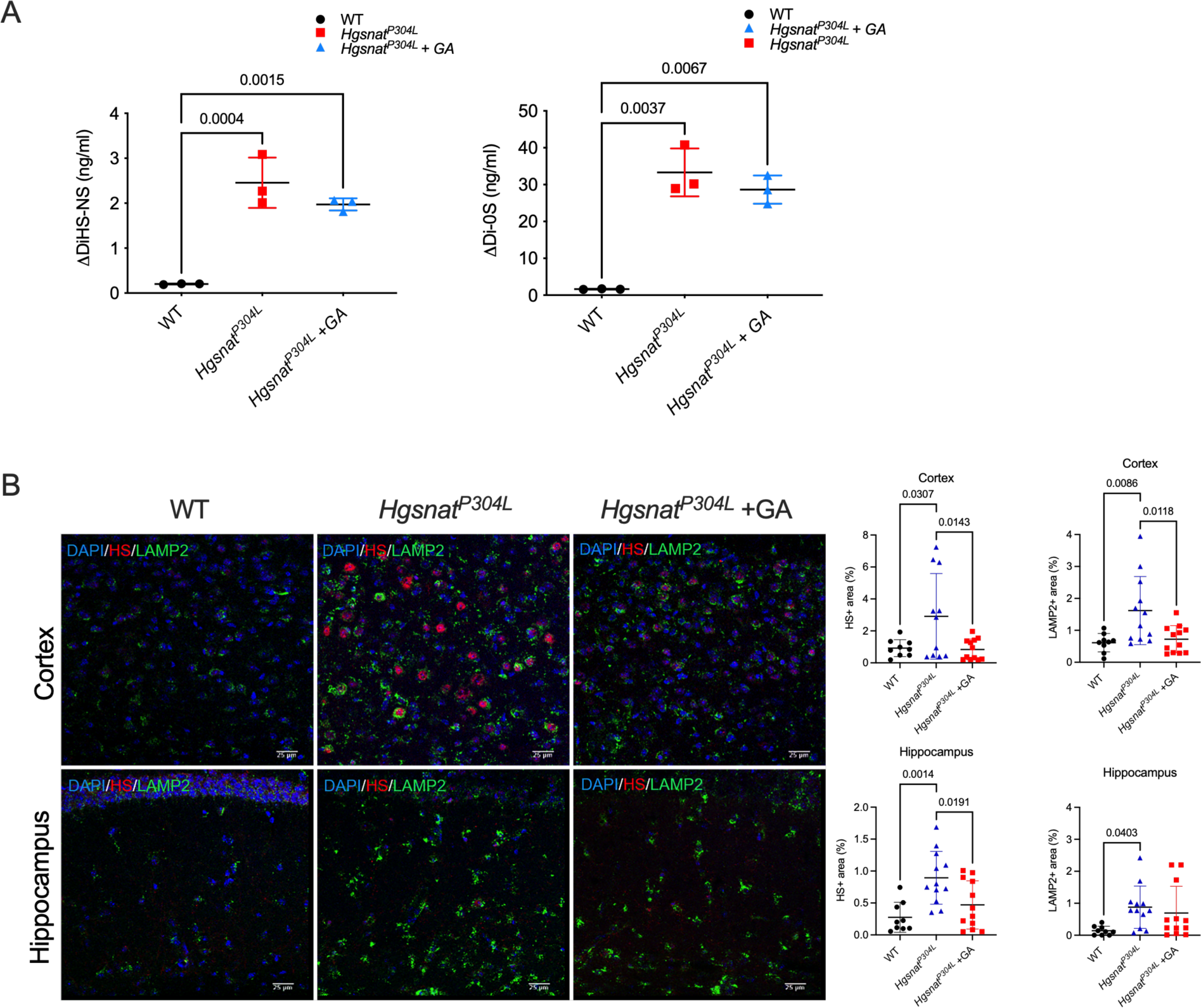
Lysosomal storage of HS is reduced in the brain of *Hgsnat^P304L^* mice treated with glucosamine. **(A)** Levels of disaccharides produced by enzymatic digestion of HS (ΔDiHS-0S and ΔDiHS-NS) measured by tandem mass spectrometry show a trend for reduction in brain tissues of 4-month- old *Hgsnat^P304L^* mice treated with glucosamine. **(B)** Reduction of HS+ and LAMP2+ puncta in cortical neurons. Panels show representative images of brain cortex (layers 4-5) and CA1 region of hippocampus of 4-month-old WT and *Hgsnat^P304L^* mice, treated or not with glucosamine, immunostained for LAMP2 (green) and HS (red). Scale bar equals 25 µm (upper and lower panels). Graphs show quantification of HS and LAMP2 staining with ImageJ software. All graphs show individual results, means and SD from experiments conducted with at least four mice (three panels per mouse) per genotype per treatment. P values were calculated using one- way ANOVA with Tukey post-hoc test.

## Discussion

Correction of a primary genetic defect either by enzyme replacement or by gene therapy/hematopoietic stem cell gene therapy remains the major approach for developing novel treatments for neurological MPS including Sanfilippo disease. However, outcomes of all recent clinical trials involving MPS III patients are either still unknown or they failed to produce desired effects, prompting researchers to explore novel alternative treatments. In particular, approaches based on the use of small molecule drugs to reduce neuroinflammation [50] (NCT04018755 clinical trial), activate autophagy [51, 52], reduce or block stored HS [53], rescue activity/expression of the mutant enzymes [8, 54], and stimulate translational readthrough of premature termination codons [55] have been described.

Development of a molecular chaperone therapy for MPS IIIC requires a generation of a novel murine model expressing misfolded HGSNAT protein. For this purpose, we selected a Pro304Leu mutation, an analog of a pathological human variant Pro311Leu, which we previously identified in severely affected MPS IIIC patients [7, 8, 40]. Now, we show that the knock-in *Hgsnat^P304L^* mouse has a drastically aggravated clinical phenotype as compared with our previously developed gene-targeted *Hgsnat-Geo* mouse model with a constitutive disruption of the gene expression [27]. In particular, the onset of the behavioral abnormalities for the knock-in *Hgsnat^P304L^* mouse was observed at 4 months, at least 2-3 months before that in the *Hgsnat-Geo* mouse. In addition, the life span was reduced by approximately 20 weeks as compared with the *Hgsnat-Geo* mouse, and, at sacrifice, the *Hgsnat^P304L^* mice showed a pronounced enlargement of kidney, liver and spleen, unlike *Hgsnat-Geo* mice that had only a mild hepatomegaly. Most of the CNS disease progression biomarkers, including neuroinflammation, astromicrogliosis, accumulation of primary and secondary storage materials, were aggravated in *Hgsnat^P304L^* mice as compared with *Hgsnat-Geo* mice of the same age. Moreover, the severity of synaptic deficits in CA1 and cultured *Hgsnat^P304L^* hippocampal neurons was increased and expanded to the GABAergic synapse. Since the residual brain level of HGSNAT acetyltransferase activity, measured with the specific BODIPY-glucosamine substrate, was reduced to below detection levels in both strains, we speculate that the difference in the clinical severity is most likely related to a toxicity of the mutant misfolded HGSNAT P304L variant expressed in the cells of the *Hgsnat^P304L^* mouse.

RNAseq analysis of mouse hippocampi demonstrated that the same types of pathways were altered in both knockout and knock-in mice as compared with their WT counterparts. In particular, lysosomal biogenesis was increased, as well as the pathways related to inflammatory and immune response. On the other hand, the expression of the genes involved in the neurogenesis and synaptic transmission was decreased. However, the number of significantly changed pathways (especially of those related to GABAergic synapse) and the degree of gene alterations were higher in the *Hgsnat^P304L^* mouse, reflecting an increased severity of the CNS pathology. Several pathways induced in the *Hgsnat^P304L^* mouse were related to the ER stress/unfolded protein response. This, together with the increased levels of the ER stress biomarkers (O-GlcNAc-modified proteins, ubiquitinated protein aggregates) detected in the hippocampal and cortical pyramidal neurons in the knock-in mouse, prompted us to hypothesize that expression of the HGSNAT mutant misfolded variant puts an additional constrain on the ER in these cells. This further impairs proteasomal protein degradation [56] and together with autophagy block, results in aggravated neuronal dysfunction, accumulation of toxic misfolded protein aggregates, neuroinflammation and eventually neurodegeneration.

To test this hypothesis, we transduced cultured hippocampal neurons established from embryos of *Hgsnat-Geo* mice with lentiviral vectors expressing either human WT HGSNAT or the P311L HGSNAT variant. In preliminary experiments, we demonstrated that the mutant HGSNAT protein was misfolded, did not have any enzymatic activity, was not proteolytically processed, did not undergo targeting to Golgi or lysosomes, and was retained in the ER. Moreover, when overexpressed, it formed cytoplasmic inclusions. When levels of synaptic proteins in cultured hippocampal *Hgsnat-Geo* neurons overexpressing WT human HGSNAT enzyme were compared to those in non-transfected cells from WT, *Hgsnat-Geo,* and *Hgsnat^P304L^* mice, we have observed a complete rescue of the markers of inhibitory and excitatory synapses. In contrast, cells expressing the mutant P311L enzyme showed a reduction in the density of inhibitory and a trend for further reduction in the density of excitatory synapses to the levels detected in the *Hgsnat^P304L^* neurons. These results directly confirm that the mutant P311L HGSNAT protein is responsible for the aggravation of synaptic deficits, especially those affecting the GABAergic synapse that are known to lead to seizures and autistic behavioural features [57].

We attempted to reduce the burden caused by the misfolded mutant enzyme in the *Hgsnat^P304L^* mice by treating them with glucosamine, a known HGSNAT chaperone. This compound increases the levels of residual HGSNAT activity in cultured fibroblasts of human MPS IIIC patients with several missense mutations, including P311L [8]. A similar effect was observed in MEF cells of *Hgsnat^P304L^* mice, confirming that the drug also has a beneficial effect on the mouse P304L HGSNAT mutant. In this study, glucosamine was provided in drinking water and was well tolerated. The treatment was started in asymptomatic mice at weaning and conducted for 13 weeks until the mice were 4-months-old. At this timepoint, behavioral testing revealed a rescue or partial rescue of defects in short-term, working, and spatial memory. Analysis of CNS pathology showed significantly reduced defects in cortical pyramidal and hippocampal CA1 neurons associated with the disease phenotype, including accumulation of autofluorescent ceroid material, primary (HS) and secondary (GM2 ganglioside, unfolded SCMAS) storage materials and induced lysosomal biogenesis. However, the treatment failed to reduce the number of astrocytes and microglia, that stayed drastically elevated in both cortex and hippocampus. It is tempting to speculate that these biomarkers, although important for assessment of overall CNS pathology progression, do not directly correlate with behavioural deficits. In contrast, the reduced levels of synaptic proteins, PSD95, VLUT1 and Syn1 in the hippocampus were rescued by glucosamine therapy once again reinforcing our previous conclusion that synaptic defects underlie the behavioural changes in MPS IIIC mice [36].

The residual HGSNAT activity in the brain and liver of treated mice showed a moderate (∼1.5 fold) but statistically significant increase suggesting a partial rescue of the mutant enzyme. Nevertheless, we cannot exclude the possibility that glucosamine induced the expression of an unknown acetyltransferase or/and a hydrolase active against unacetylated Muf-*β*-glucosaminide substrate. Although, we have observed a promising behavioral improvement, it remains to be determined whether such small increase in the residual HGSNAT activity and HS catabolism is sufficient to produce a long-term correction of the clinical phenotype in *Hgsnat^P304L^* mice. Alternatively, glucosamine can reduce the pool of the misfolded HGSNAT variant accumulating in the ER, thus decreasing the load of ubiquitin-mediated protein degradation pathway and ER stress. Paradoxically, glucosamine which is known to induce the ER stress and unfolded protein response in multiple types of cells (reviewed in [58]), did not induce ER stress markers in the brain of treated mice, including accumulation of ubiquitinated aggregates, or levels of O-GlcNAc-modified proteins, which at 4 months remained at the level of WT mice in both treated and untreated groups.

Administration of glucosamine in concentrations of 1-2 g/kg/day similar to those we used in this study is safe in mice, but in humans it is hardly achievable[59]. Thus, identifying HGSNAT chaperones with higher potency and bioavailability either by library screening or a rational design is required before such an approach can be tested in human patients. Besides, since glucosamine does not increase residual activity of the deficient N-sulfoglucosamine sulfohydrolase (SGSH) enzyme in the cultured fibroblasts of MPS IIIA patients affected with missense mutations (Martins and Pshezhetsky, unpublished), specific chaperones need to be identified for other subtypes of MPS III. Importantly, our current study established that, while secondary accumulation of misfolded proteins and gangliosides showed a drastic response to chaperone treatment, neuroinflammation was not reduced. This suggests that the efficacy of chaperone therapy can be further improved when combined with drugs that primarily address neuroinflammation such as recently described blockers of the IL-1*β* pathways [50].

In conclusion, the results of this study validate the novel *Hgsnat^P304L^* mouse strain as a model closely mimicking the phenotype of the early onset, fast progressing MPS IIIC patients, useful for testing prognostic biomarkers of the disease and novel therapies. Importantly, our work establishes that misfolded mutant HGSNAT protein expression is a key contributing factor for the pathophysiology of MPS IIIC and suggests that treatment with chaperones, capable of rescuing the folding process of such mutants, should be considered as an alternative therapy for this disease.

## Availability of data and materials

All data generated or analyzed during this study are included in this published article and its supplementary information files.

## Authors’ contributions

Conducted experiments and acquired data: X.P., M.T.; P.B., R.H-R., A.L.A.N., TM.X., C.P., D.P. and S.K.; analyzed data: X.P., M.T.; P.B., R.H-R., A.L.A.N., TM.X., D.P., S.K., S.T., F.M.P., C.R.M., and A.V.P.; wrote the manuscript (first draft) X.P., M.T., P.B., A.V.P.; wrote the manuscript (editing) A.V.P., S.T., F.M.P. and C.R.M. All authors read and approved the final manuscript.

## Acknowledgements

The authors thank Dr. Jeffrey A. Medin, Dr. Monty McKillop, Dr. Christian Beauséjour and Gaël Moquin-Beaudry for the help in production of LV-HGSNAT-GFP and LV-P311L- HGSNAT-GFP. We also thank Jeannie Mui and the Facility for Electron Microscopy Research (FEMR – McGill University) for the help with the transmission electron microscopy, Dr. Elke Küster-Schöck and the Plateforme d’Imagerie Microscopique (PIM – CHU Sainte Justine) for the help with life imaging microscopy, Mitra Cowan and the McGill Integrated Core for Animal Modeling (MICAM) for the help with mouse production and Dr. Mila Ashmarina for critically reading the manuscript and helpful advice.

