## Supplemental Figures and Tables for "Oral Glucosamine Ameliorates Aggravated Neurological Phenotype in Mucopolysaccharidosis III Type C Mouse Model Expressing Misfolded HGSNAT Variant"

### Supplementary Data

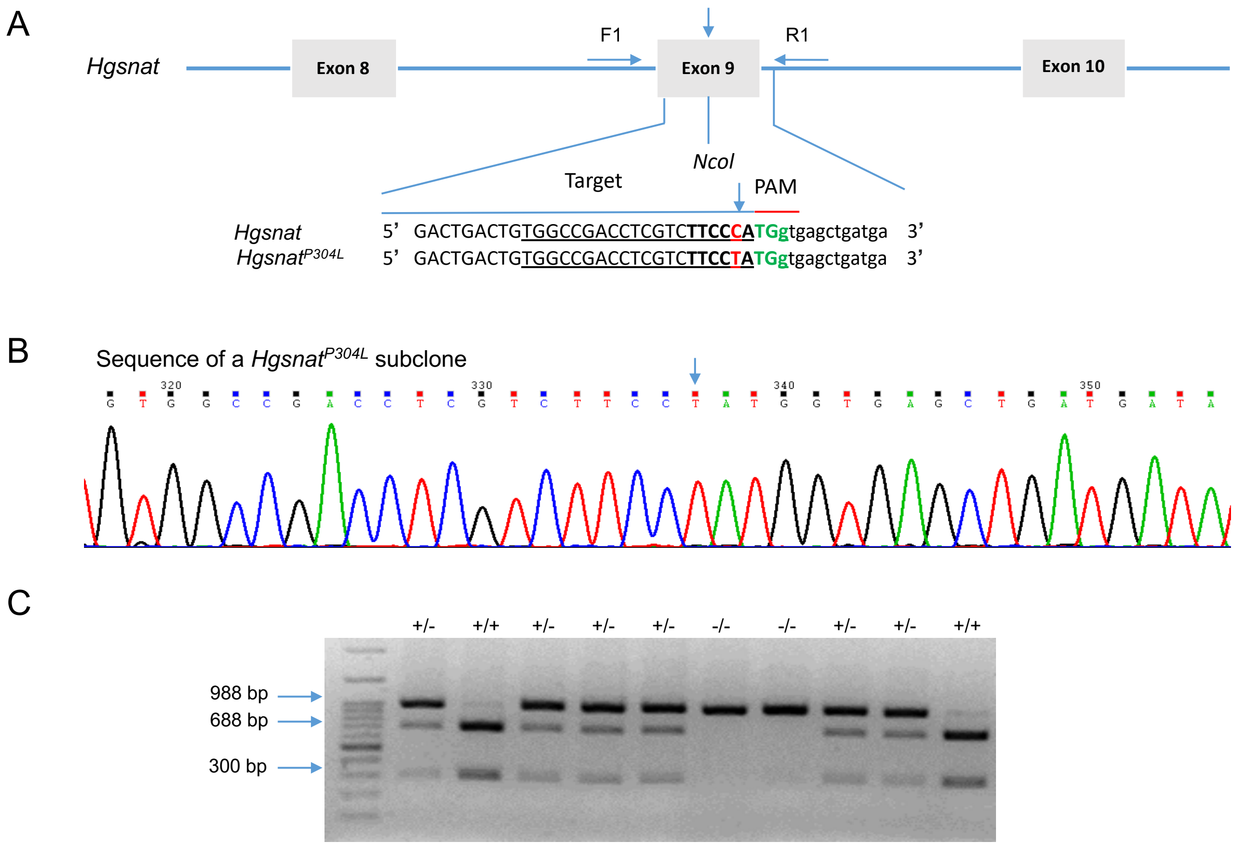


**Figure S1. Generation and genotyping of *Hgsnat*$^{P304L}$ mice.**

**(A)** Schema showing the Cas9/sgRNA-targeting site in *Hgsnat* exon 9. The sgRNA-targeting sequence is underlined, and the protospacer-adjacent motif (PAM) sequence is shown in green. The c.911C>T mutation is shown in red and marked with an arrow. The C>T substitution disrupts the *NcoI* restriction site (shown in bold). The exon sequence is capitalized. **(B)** Sanger sequencing of single allele fragment, obtained by PCR amplification of genomic DNA from the tail clips of the *Hgsnat*$^{P304L}$ founder mouse showing the presence of the c.911C>T mutation. **(C)** Genotyping of *Hgsnat*$^{P304L}$ mice. The DNA was extracted from clipped mouse tails and a 988-bp product amplified using a forward primer ATGGAGTGCCTGATGGGAGG and a reverse primer GATCTAGAAACGGCCCGAAGA. The PCR products were further digested with *NcoI* and analyzed on a 2% agarose gel. The 688 and 300-bp fragments are detected for the WT allele, and an undigested 988-bp fragment for the targeted *Hgsnat*$^{P304L}$ allele.

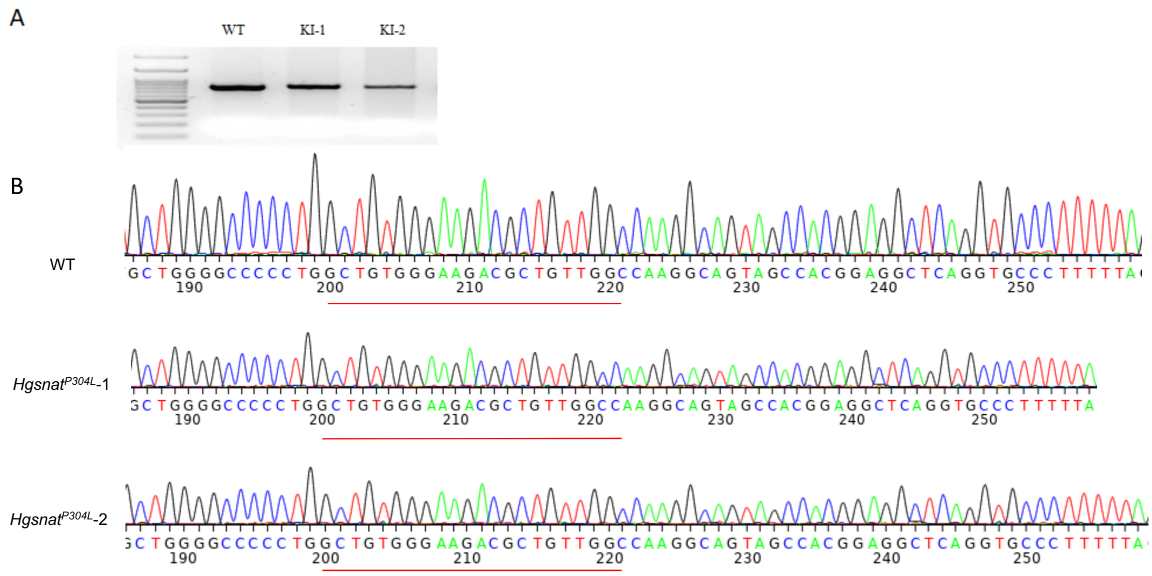


**Figure S2. Absence of off-target effects in the *Hgsnat*$^{P304L}$ founder mice.**

**(A)** A 763-bp fragment of the *Spg7* gene containing the potential off-target sequence CTGTGGGAAGACGCTGTTGGCCA was amplified by PCR from DNA extracted from tail clips of *Hgsnat*$^{P304L}$ founder mice (KI-1 and KI-2) and a control WT mouse. **(B)** Sanger sequencing of a PCR product confirms the absence of mutations in the *Spg7* gene fragment adjacent to the CTGTGGGAAGACGCTGTTGGCCA fragment homologous to the PAM sequence.

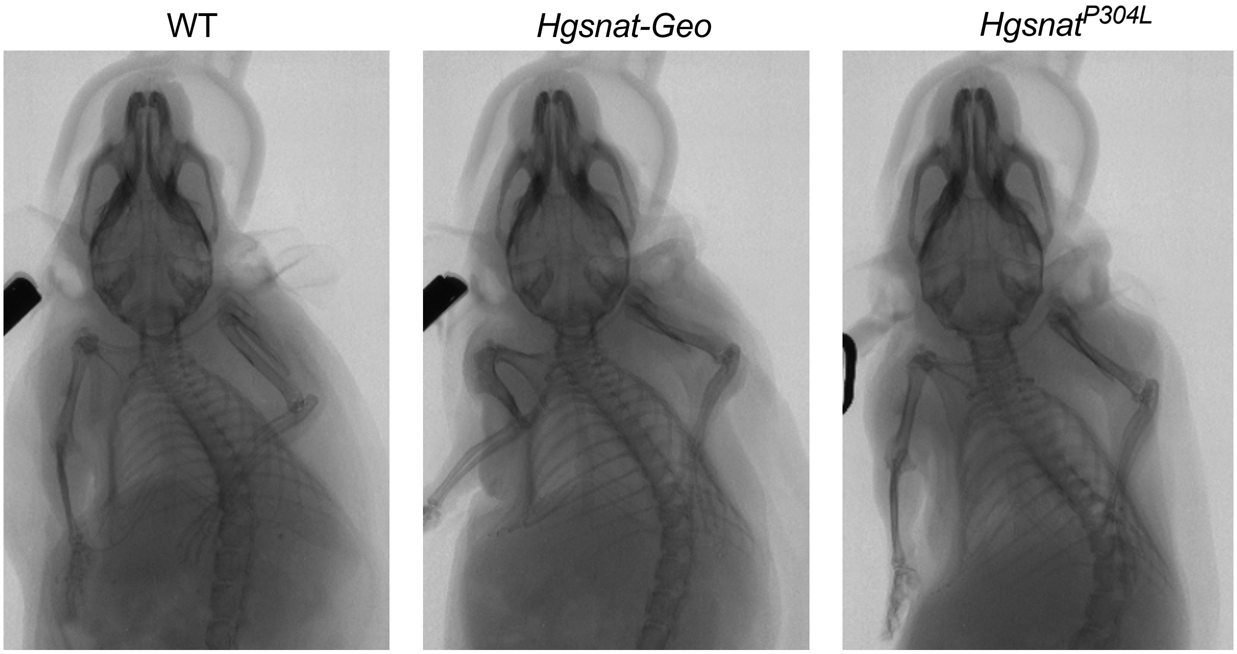


**Figure S3. *Hgsnat-Geo* and *Hgsnat*$^{P304L}$ mice do not develop abnormalities of skull bones.** A high-resolution in vivo micro-CT scanner (SkyScan 1176) was used to evaluate skeletal deformities in 4-months-old *Hgsnat-Geo* and *Hgsnat*$^{P304L}$ mice. The mice were anesthetized by isoflurane flow and the images were taken from the dorsal side.

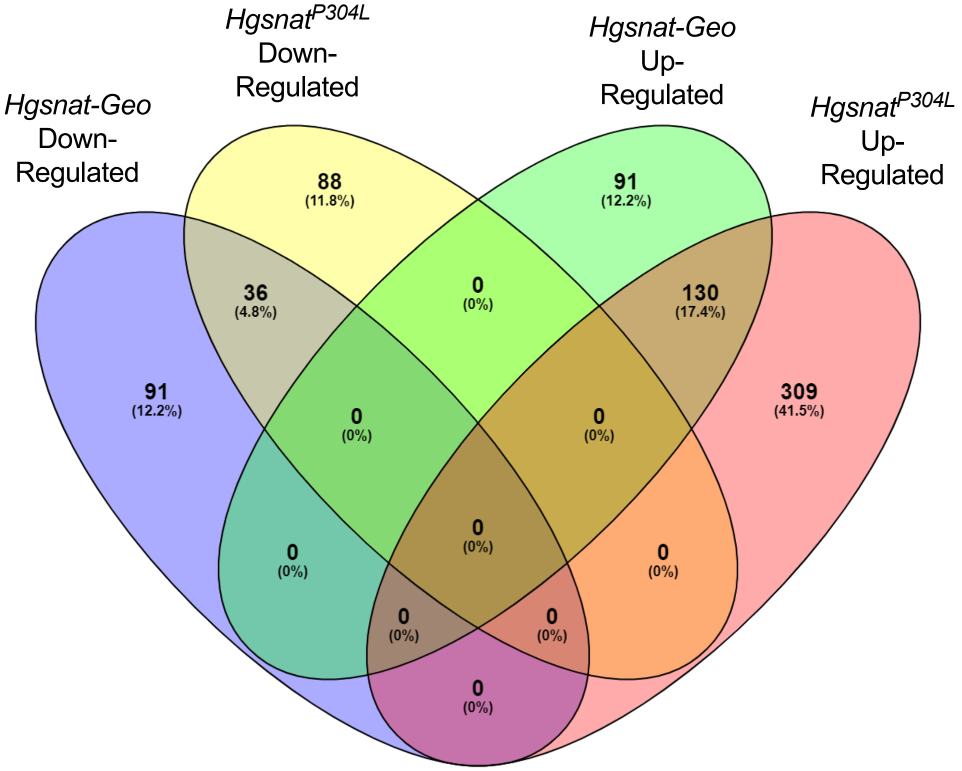


**Figure S4. A higher number of hippocampal genes with altered expression levels as compared with WT mice is found in *Hgsnat*$^{P304L}$ than *Hgsnat-Geo* mice.**

Venn diagram showing the number of genes that were up-regulated or down-regulated in hippocampal tissues of 4-month-old *Hgsnat*$^{P304L}$ and *Hgsnat-Geo* as compared with the age- and sex-matched WT mice. Three mice (two males and one female) were studied for each genotype.

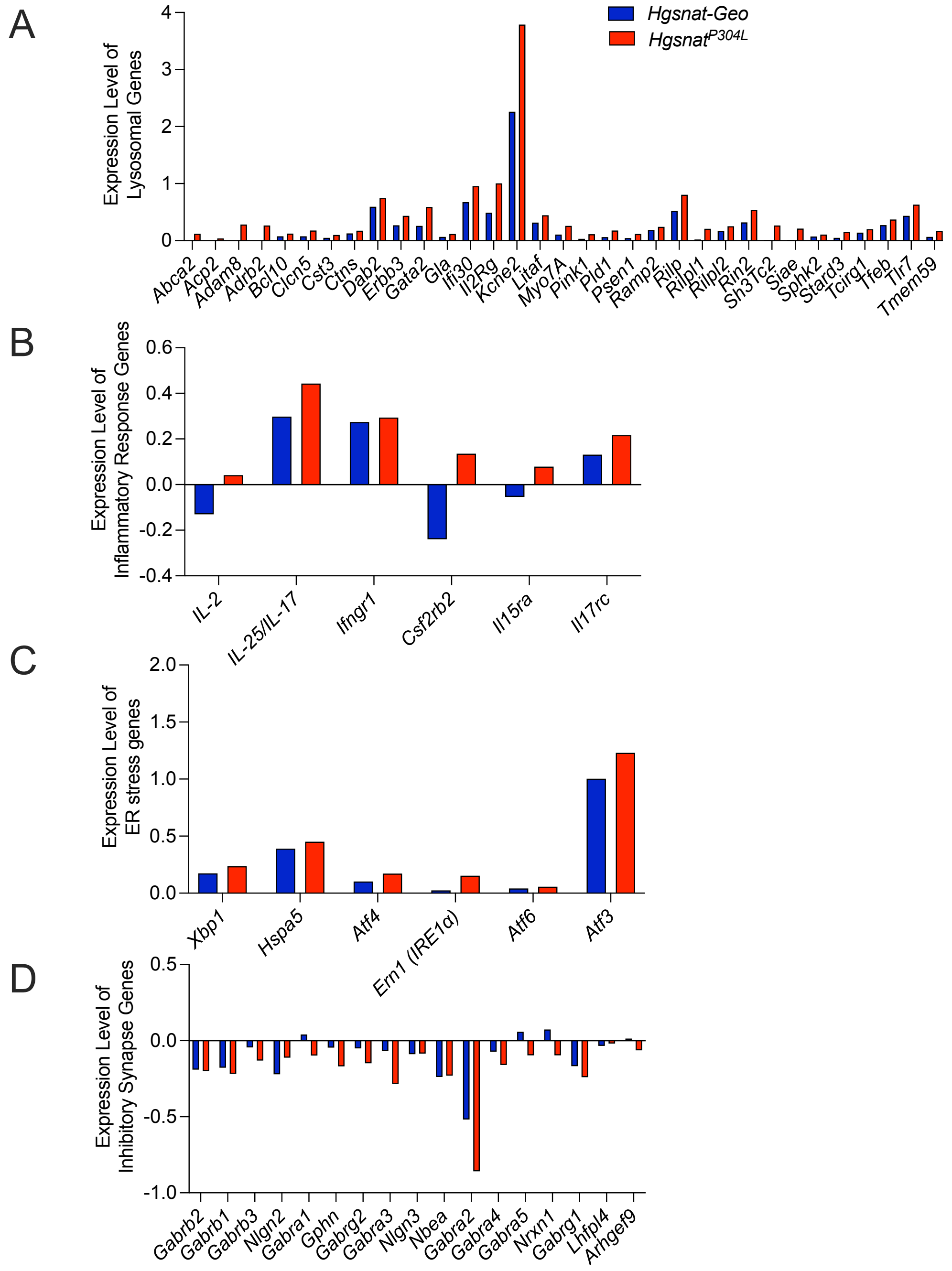


**Figure S5. Expression levels of genes involved in lysosomal biogenesis, inflammatory response, ER stress and inhibitory synapse show a trend for increased change in *Hgsnat*$^{P304L}$ as compared with *Hgsnat-Geo* mice.**

The expression levels of genes involved in lysosomal biogenesis **(A)**, inflammation **(B)**, and UPR **(C)** show a trend for increase, while the expression of genes involved in inhibitory synapse **(D)** show a trend for decrease in *Hgsnat*$^{P304L}$ as compared with *Hgsnat-Geo* mice.

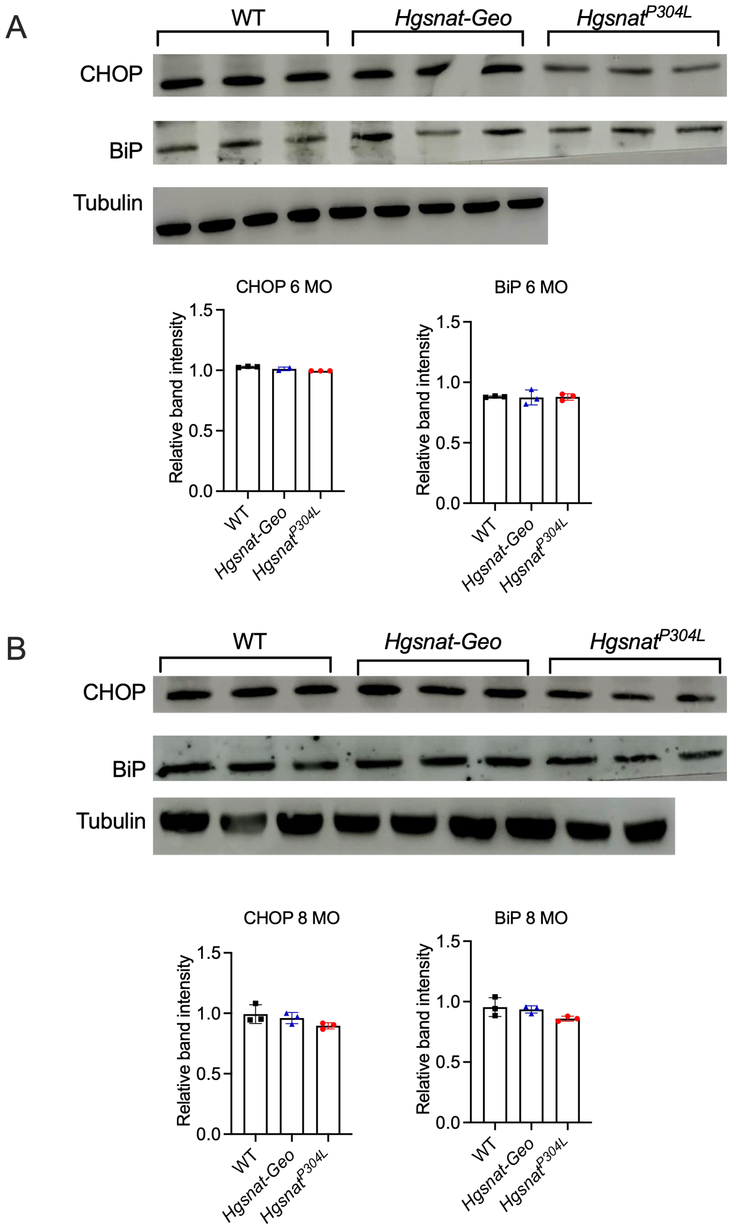


**Figure S6. Normal protein level of ER stress markers CHOP and BiP are found in brain cortex tissues of both *Hgsnat*$^{P304L}$ and *Hgsnat-Geo* mice at the age of 6 and 8 months.**

Representative immunoblots showing expression levels of ER stress markers in the cortices of 6-month-old **(A)** and 8-month-old **(B)** WT, *Hgsnat*$^{P304L}$, and *Hgsnat-Geo mice*. Graphs show band intensity values measured using ImageJ software. Individual results, means, and SD of experiments with 3 mice per genotype, per age are shown. P values were calculated using one-way ANOVA with Tukey post-hoc test.

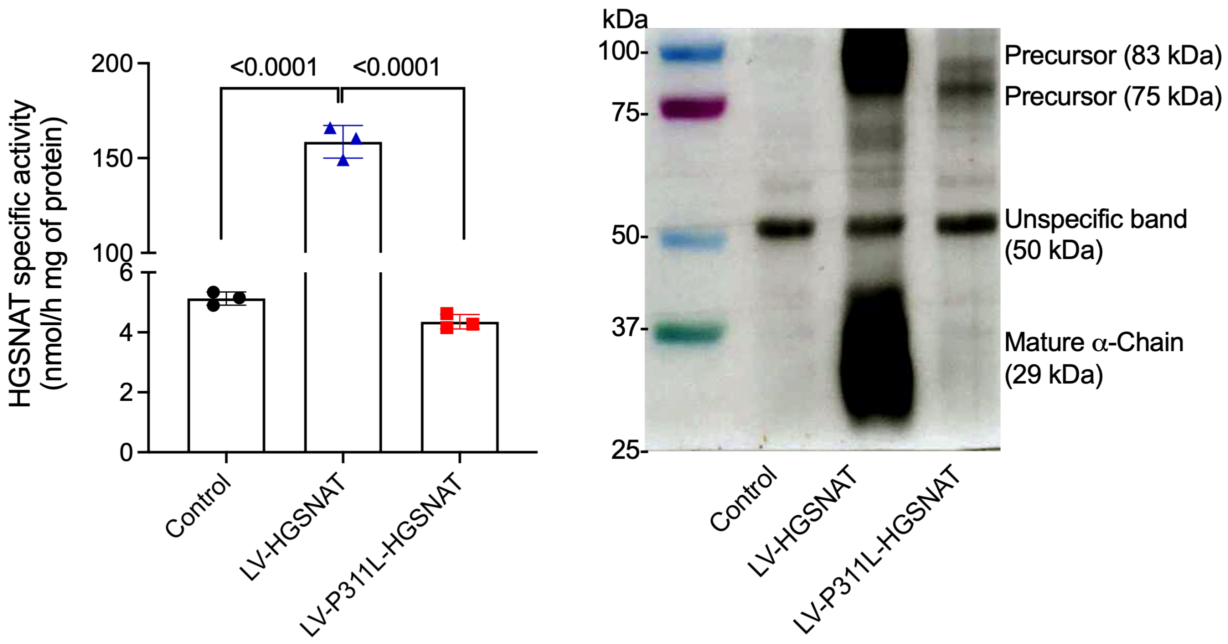


**Figure S7. The missense variant Pro311Leu affects expression, processing and enzymatic activity of HGSNAT.**

**(A)** Pro311Leu HGSNAT mutant lacks enzymatic activity. The N-acetyltransferase activity was essayed in homogenates of cultured skin fibroblasts of healthy control donor (Control) transduced with LV vectors encoding for the GFP-tagged WT HGSNAT (LV-HGSNAT) and or the Pro311Leu mutant (LV-P311L-HGSNAT). The graph shows individual values, means and SD of three independent experiments. *P*-values were calculated by one-way ANOVA followed by Tukey's post-hoc multiple comparison test. **(B)** The 75 kDa (with EGFP tag) non-glycosylated precursor is the main form detected in the homogenates of cells transduced with the mutant virus, while the fully glycosylated 83 kDa precursor and the cleaved 29 kDa α-subunit are detected in cells expressing the WT enzyme. The 50 kDa band represents a non-specific cross-reacting protein also present in non-transduced cells. The panel shows a representative blot of two independent experiments yielding similar results.

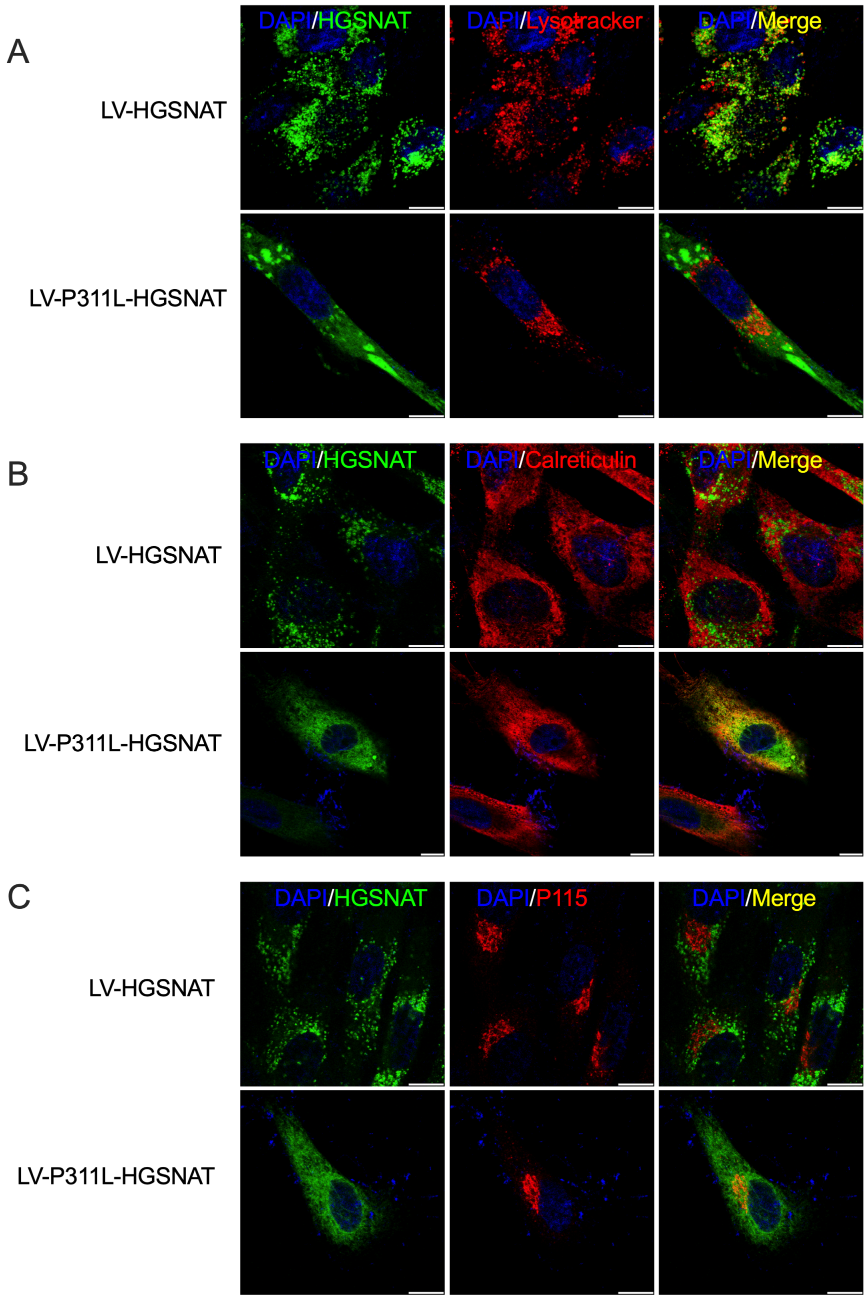


**Figure S8. The Pro311Leu HGSNAT mutant protein is not targeted to lysosomes.**

Representative confocal images show fibroblast cells transduced with LV vectors encoding for the GFP-tagged WT HGSNAT (LV-HGSNAT) and or the Pro311Leu mutant (LV-P311L-HGSNAT). Cells grown on glass slides were labeled with Lysotracker Red for one hour before fixation **(A)** or immunostained for the ER (anti-Calreticulin antibodies) **(B)** or Golgi (anti-P115 antibodies) **(C)** (red). Scale bar: 10 μm.

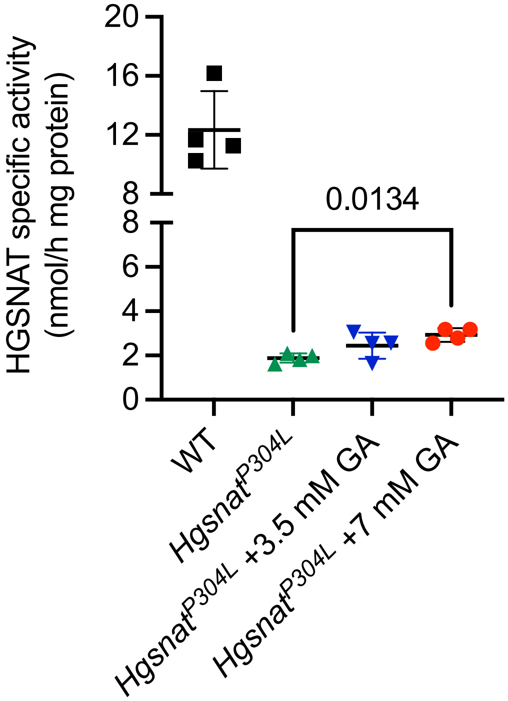


**Figure S9. Glucosamine significantly increases HGSNAT activity in cultured mouse embryonic fibroblasts (MEF cells) of homozygous *Hgsnat*$^{P304L}$ mice.**

HGSNAT activity was measured with fluorogenic substrate, Muf-β-D-glucosaminide in homogenates of cultured mouse embryonic fibroblasts (MEF cells) of homozygous *Hgsnat*$^{P304L}$ and WT mice treated or not with glucosamine for 5 days. Graph shows individual results, means and SD of experiments conducted with four different cell cultures. P values were calculated using one-way ANOVA with Tukey post-hoc test.

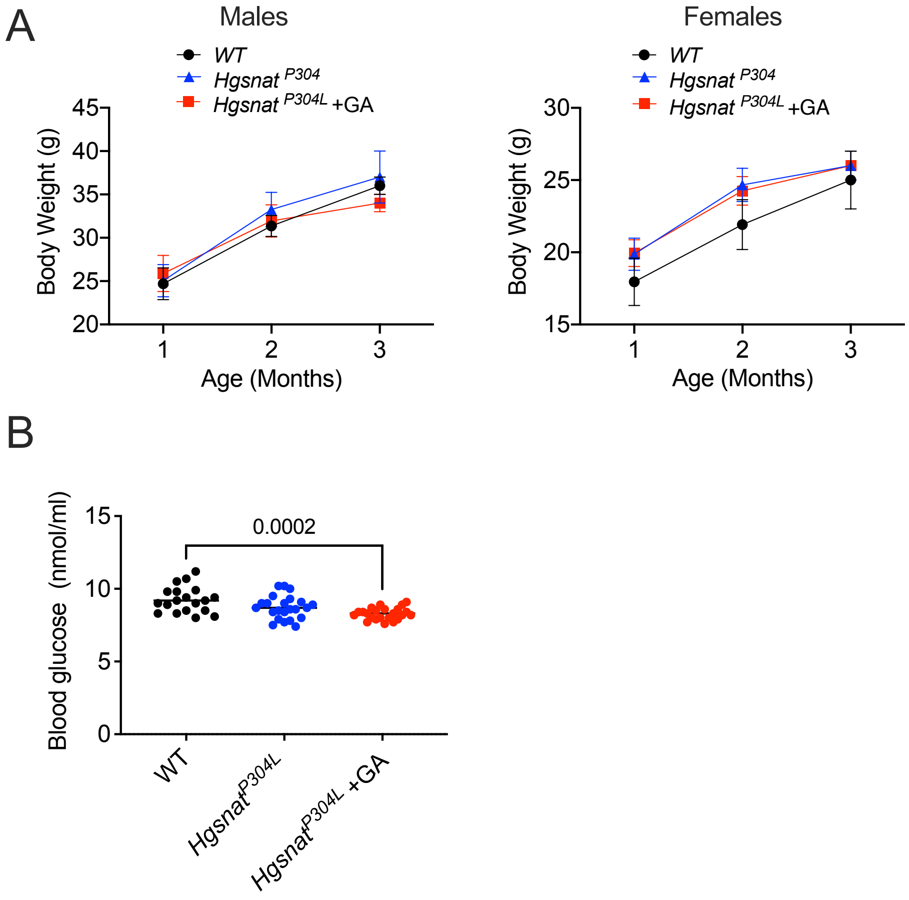


**Figure S10. Mice treated daily with 2.0 g/kg BW of glucosamine for 13 weeks do not show alterations in growth and body weight or increase in blood glucose levels.**
**(A)** Mice treated daily with 2.0 g/kg BW of glucosamine for 13 weeks do not show alterations in growth. Body weight was measured monthly, between the age of one and three months. Mean values and SD obtained for 12 mice per genotype, per gender, per treatment are shown. **(B)** The blood glucose levels were tested at the age of 4 months. Individual data, means and SD obtained for at least 24 mice per genotype, per treatment are shown. P values were measured using two-way ANOVA **(A)** and one-way ANOVA **(B)** with Tukey post-hoc tests.

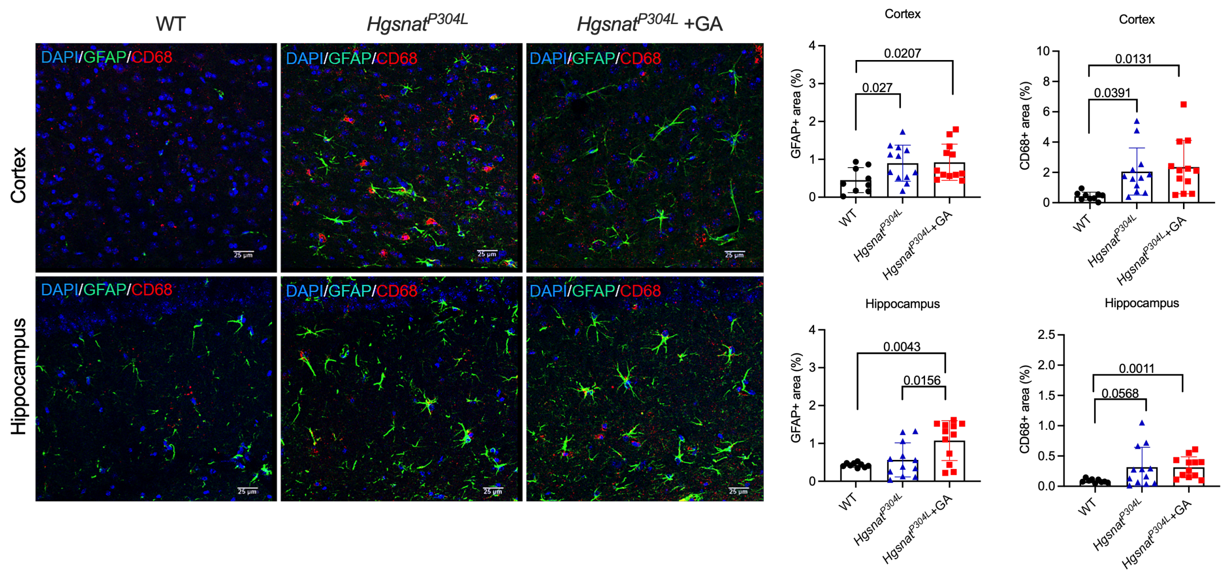


**Figure S11. The levels of activated CD68+ microglia and GFAP+ macrophages are not changed in the hippocampus and somatosensory cortex of 4-month-old *Hgsnat*$^{P304L}$ mice treated with glucosamine.**

Panels show representative images of brain cortex (layers 4-5) and hippocampus of 4-month-old WT, and *Hgsnat*$^{P304L}$ mice treated or not with glucosamine and immunostained for GFAP (green) and CD68 (red). Scale bars equal 25 µm. Bar graph shows quantification of CD68+ and GFAP+ area with ImageJ software. Individual results, means, and SD of experiments with four mice per genotype, per treatment are shown. P values were calculated using one-way ANOVA with Tukey post-hoc test.

**Table S1. Top ten upregulated and downregulated differentially expressed genes in *Hgsnat-Geo* mice as compare with WT mice. All results are available for download and analysis at the following address:**

**WT/KI:**
**https://maayanlab.cloud/biojupies/notebook/IAykq58fQ**

**WT/KO:**
**https://maayanlab.cloud/biojupies/notebook/m2KTE2Z4W**

**KO/KI:**
**https://maayanlab.cloud/biojupies/notebook/8Hq6GUKin**

1.1 The top 10 upregulated differentially expressed genes

| Gene Symbol | logFC | AveExpr | t | P-value | Adj.P.Val | B |
|---|---|---|---|---|---|---|
| Gm15446 | 3.963596 | 3.298196 | 25.95803 | 5.09E-09 | 8.00E-05 | 2.83828 |
| Lyz2 | 3.293204 | 3.47975 | 11.79959 | 2.40E-06 | 0.018907 | 2.05529 |
| Tmem254a | 1.637335 | 3.025582 | 10.69658 | 5.06E-06 | 0.019894 | 1.423174 |
| Tmem254b | 1.637335 | 3.025582 | 10.69658 | 5.06E-06 | 0.019894 | 1.423174 |
| Ly9 | 3.165477 | -0.32953 | 9.831997 | 9.52E-06 | 0.029949 | -1.91727 |
| Mpeg1 | 1.380629 | 5.344022 | 8.07209 | 4.06E-05 | 0.106301 | 1.835164 |
| Lilrb4a | 2.845683 | -1.42679 | 7.39661 | 7.58E-05 | 0.170301 | -2.79881 |
| Cst7 | 3.299472 | -0.71795 | 6.989179 | 0.000113 | 0.220437 | -2.45246 |
| Clec7a | 2.958558 | 0.58566 | 6.803532 | 0.000136 | 0.220437 | -1.60498 |
| Cd68 | 1.33784 | 3.060377 | 6.776075 | 0.00014 | 0.220437 | 0.173089 |

1.2 The top 10 downregulated differentially expressed genes

| Gene Symbol | logFC | AveExpr | t | P-value | Adj.P.Val | B |
|---|---|---|---|---|---|---|
| Tnfrsf18 | -1.02505 | 1.123303 | -4.41071 | 0.002245 | 0.965579 | -2.15811 |
| Tgfb1i1 | -0.72034 | 3.124157 | -4.25156 | 0.002783 | 0.965579 | -1.55517 |
| Gm16432 | -0.82374 | 2.677928 | -4.06556 | 0.003593 | 0.98568 | -1.83306 |
| Tmem91 | -0.61836 | 2.40964 | -3.97921 | 0.004053 | 0.98568 | -1.98018 |
| Col2a1 | -1.12817 | -0.09692 | -3.94625 | 0.004245 | 0.98568 | -2.93069 |
| Cxcl15 | -0.81951 | 1.966697 | -3.76807 | 0.005465 | 0.998245 | -2.28399 |
| Phldb3 | -1.32778 | -1.23601 | -3.65429 | 0.006436 | 0.998245 | -3.43336 |
| Sptssb | -1.05821 | 0.310449 | -3.63897 | 0.006581 | 0.998245 | -2.93564 |
| Gdpd2 | -0.66009 | 2.526844 | -3.63574 | 0.006612 | 0.998245 | -2.2641 |
| Mcm10 | -1.07276 | 0.078383 | -3.63064 | 0.006661 | 0.998245 | -3.021 |

**Table S2. Top ten upregulated and downregulated differentially expressed genes in *Hgsnat*$^{P304L}$ mice as compared with WT mice.**

2.1 The top 10 upregulated differentially expressed genes

| Gene Symbol | logFC | AveExpr | t | P-value | Adj.P.Val | B |
|---|---|---|---|---|---|---|
| Gm15446 | 3.863174 | 3.274976 | 31.89635 | 5.80E-11 | 9.21E-07 | 11.93097 |
| Zfp781 | 6.678025 | -2.12467 | 15.55362 | 4.61E-08 | 0.000367 | 2.134689 |
| Gm3095 | 4.992654 | -2.95406 | 10.01197 | 2.39E-06 | 0.012683 | 0.616014 |
| Tmem254b | 1.436813 | 2.935904 | 9.158899 | 5.18E-06 | 0.015925 | 4.51273 |
| Tmem254a | 1.436813 | 2.935904 | 9.158899 | 5.18E-06 | 0.015925 | 4.51273 |
| Sema3b | 2.028503 | 1.460513 | 9.001059 | 6.01E-06 | 0.015925 | 3.850118 |
| Trpv4 | 2.375905 | 1.397375 | 8.776806 | 7.46E-06 | 0.01625 | 3.660536 |
| Trf | 0.703259 | 7.395393 | 8.682262 | 8.18E-06 | 0.01625 | 4.148475 |
| Mobp | 0.779389 | 9.230635 | 8.448458 | 1.03E-05 | 0.018209 | 3.909972 |
| Car14 | 1.312602 | 2.812631 | 7.765717 | 2.09E-05 | 0.033208 | 3.239745 |

2.2 The top 10 downregulated differentially expressed genes

| Gene Symbol | logFC | AveExpr | t | P-value | Adj.P.Val | B |
|---|---|---|---|---|---|---|
| Plxnd1 | -0.53093 | 4.796305 | -5.33373 | 0.000397 | 0.100643 | 0.233987 |
| Oprk1 | -1.10921 | 0.966172 | -5.006 | 0.000627 | 0.129846 | -0.01527 |
| Dcx | -0.52407 | 4.870583 | -4.81783 | 0.000821 | 0.137388 | -0.52297 |
| Gm5884 | -3.88558 | -0.98275 | -4.53303 | 0.001246 | 0.170947 | -0.94441 |
| Tmem181b-ps | -1.02276 | 6.272376 | -4.48815 | 0.001332 | 0.173558 | -1.11689 |
| Tmem181a | -0.70504 | 8.02711 | -4.31821 | 0.001719 | 0.210769 | -1.39994 |
| Gm3173 | -3.71812 | -2.42109 | -4.27973 | 0.001823 | 0.21311 | -1.60701 |
| Cntnap3 | -0.93077 | 2.250784 | -4.18608 | 0.002103 | 0.219739 | -1.12029 |
| Pde11a | -0.76544 | 4.031976 | -4.15494 | 0.002206 | 0.22343 | -1.41855 |
| Gm14253 | -1.9134 | 0.027017 | -3.95317 | 0.003017 | 0.255198 | -1.43675 |

**Table S3. Top ten upregulated and downregulated differentially expressed genes in *Hgsnat*$^{P304L}$ mice as compared with *Hgsnat-Geo* mice.**

3.1 The top 10 upregulated differentially expressed genes

| Gene Symbol | logFC | AveExpr | t | P-value | Adj.P.Val | B |
|---|---|---|---|---|---|---|
| Gm42421 | 4.052645 | -2.63085 | 5.214733 | 0.000774 | 0.999853 | -4.52378 |
| Gm3095 | 3.670507 | -2.34643 | 4.229227 | 0.002795 | 0.999853 | -4.52615 |
| Xkr7 | 1.708508 | -1.3427 | 3.981301 | 0.003947 | 0.999853 | -4.49992 |
| Foxb1 | 1.423888 | -1.0626 | 3.518206 | 0.007702 | 0.999853 | -4.50034 |
| Gm13543 | 1.678381 | -1.66592 | 3.356957 | 0.009787 | 0.999853 | -4.52317 |
| Scnn1a | 0.537885 | 2.606791 | 3.285439 | 0.010896 | 0.999853 | -4.20182 |
| Gm13369 | 2.098103 | -1.5228 | 3.263089 | 0.011269 | 0.999853 | -4.52148 |

| | | | | | | |
|---|---|---|---|---|---|---|
| 5031410I06Rik | 4.496545 | -1.07879 | 3.240814 | 0.011654 | 0.999853 | -4.50863 |
| Lrrc43 | 1.200727 | -1.02501 | 3.22557 | 0.011925 | 0.999853 | -4.50728 |
| Pcsk4 | 0.928758 | 0.128132 | 3.209835 | 0.012212 | 0.999853 | -4.45661 |

3.2 The top 10 downregulated differentially expressed genes

| Gene Symbol | logFC | AveExpr | t | P-value | Adj.P.Val | B |
|---|---|---|---|---|---|---|
| Rab11b-ps2 | -2.66777 | 0.558057 | -5.55964 | 0.00051 | 0.999853 | -4.33989 |
| Gm5884 | -3.6254 | -1.07655 | -3.67126 | 0.006155 | 0.999853 | -4.49694 |
| Duxbl1 | -4.11426 | -1.6153 | -3.52989 | 0.007571 | 0.999853 | -4.51799 |
| Btg3 | -0.61514 | 3.991453 | -3.32243 | 0.010307 | 0.999853 | -3.97252 |
| Ddx3y | -4.95021 | 3.824088 | -2.93646 | 0.018542 | 0.999853 | -4.0868 |
| Lilrb4a | -1.34239 | -0.7047 | -2.80853 | 0.022597 | 0.999853 | -4.51101 |
| Eif2s3y | -5.06998 | 3.206903 | -2.79523 | 0.023069 | 0.999853 | -4.20187 |
| Nmrk2 | -0.89672 | -0.23561 | -2.79509 | 0.023074 | 0.999853 | -4.49442 |
| Gm10357 | -1.60994 | 0.746493 | -2.78907 | 0.02329 | 0.999853 | -4.44375 |
| Gm28064 | -1.01025 | -0.79637 | -2.75859 | 0.02442 | 0.999853 | -4.5159 |
